# Non-linearity of spatial integration varies across layers of primary visual cortex

**DOI:** 10.1101/2025.04.10.648107

**Authors:** Remy Cagnol, Ján Antolík, Larry A Palmer, Diego Contreras

## Abstract

The receptive field (RF) of visual cortical neurons is highly dynamic and context-dependent, shaped by both the spatial and temporal properties of stimuli and the complex architecture of cortical circuits. While classical RF mapping through extracellular recordings reveals only the area triggering spiking responses, intracellular recordings reveal a much broader region of subthreshold synaptic input. We investigated how neurons in different cortical layers integrate visual input across space, with a focus on the linearity of spatial summation. Using intracellular recordings, we found that supragranular complex cells integrate input in a highly sublinear manner, in contrast to infragranular complex cells and simple cells, which exhibited near-linear summation. To understand the underlying mechanisms, we employed a large-scale recurrent spiking model of cat primary visual cortex (V1). Modeling results point to the differential patterning of long-range horizontal connections—particularly their targeting of excitatory versus inhibitory neurons—as a potential source of the observed layer-specific integration properties. These findings suggest that RFs emerge from interaction of feedforward, horizontal, and possibly feedback inputs, that are continuous in space, challenging the conventional notions of fixed spatial RF boundaries in early visual processing.

- *The properties of receptive fields (RF) of neurons in the primary visual cortex (V1) are not static but change depending on spatiotemporal properties of visual stimulus*.
- *To study spatial integration of visual information by V1 neurons in a RF independent way, we made intracellular recordings across the cat V1 in response to drifting gratings confined to either circular apertures of variable diameters (disks), or in annuli with fixed outer diameters and variable inner diameters (rings)*.
- *We find that supragranular complex cells integrate input in a sublinear manner, whereas infragranular complex cells and simple cells exhibit near-linear summation*.
- *Our large-scale recurrent spiking model shows that these differences between supragranular and infragranular complex cells can be explained by laminar differences in horizontal connectivity properties*.
- *These results highlight the substantial differences in spatial integration of visual information at different stages of visual processing*

## Introduction

The primary function of neurons in the visual cortex is to integrate visual information over space and time. The receptive field (RF) of a neuron is defined as the area of visual space where stimuli modulate its response (Hartline, 1938) (Kuffler, 1953; Hubel and Wiesel, 1959). While often treated as immutable, the properties of the RF strongly depend on the measurement technique. Extracellular recordings of spikes produce RF maps of only those visual inputs that reach firing threshold (usually called the classical receptive field (Barlow, 1953)), while intracellular recordings reveal a much larger area of visual space from which visual stimuli trigger only subthreshold depolarisation, thus generating a much larger RF map (Bringuier et al., 1999a; Angelucci and Shushruth, 2013). Furthermore, if the spiking or classical RF is measured by probing space with small stimuli (the minimum discharge field of Barlow et al. (1967)), they appear much smaller than if using patches of drifting gratings of increasing area (DeAngelis et al., 1994; Sceniak et al., 1999; Cavanaugh et al., 2002). Shortly after the establishment of the concept of RF it was found that stimuli placed in its proximity, that cannot drive the neural response on its own, can however modulate the neuron’s response to a stimulus placed inside the RF. These modulatory effects were most often found to be suppressive, giving rise to the concept of suppressive surround. However, this artificial “boundary” between a center and a surround is not fixed, depending on range of stimulus properties, most notably its contrast and its orientation with respect to the center (Gilbert et al., 1996; Kapadia et al., 1999; Sceniak et al., 1999; Angelucci et al., 2017).

Rather than constrained by artificial boundaries, or acting as a set of interacting discrete regions, RFs emerge from computations performed by the underlying neural substrates which unfold in continuous space and time. Synaptic inputs arising from a multiplicity of sources contribute to the emergence of RFs. Retinal output is relayed to cortex by lateral geniculate axons arriving in layers 4 and 6 (Binzegger et al., 2004; Stepanyants et al., 2008). There, they activate local circuits of excitatory and inhibitory neurons which in turn propagate activity vertically through the functional column (Peters and Payne, 1993). Through a system of long-range horizontal unmyelinated axons terminating in discrete patches in supra and infragranular layers, columnar activation then spreads away from the retinotopic activation site to a large area of surrounding cortex. Finally, the distribution of activation by visual input rapidly engages higher visual areas which generate feedback to upper layers of primary visual cortex (V1)[r]. In turn, activation of layer 6 neurons generates feedback to the same LGN cells which are the source of the feedforward input[r]. Crucially, both the horizontal and feedback pathway form recurrent networks that endow the system with complex dynamics that ultimately give rise to what is measured as the receptive field. Despite extensive knowledge of each one of these elements on the complex array of circuits and cells involved in visual representation at the early stages of visual processing, we are very far from understanding the dynamics of the system in response to naturalistic scenes succeeding rapidly in time during natural vision.

Thus, the synthesis of RFs is a combination of multiple recurrent circuits, only a small percent of which are activated by feedforward thalamic input. The vast majority of inputs are cortically generated, including intralaminar, interlaminar, horizontal and feedback connections. The complex dynamics of activation of these circuits is dependent on stimulus properties and the state of activation of the network and thus, strict spatial definitions of RFs are either simply semantic or operational for the purpose of a particular experimental design.

A question of fundamental importance to understand the computations performed by visual cortex is the degree of linearity of spatial summation of visually-driven synaptic input by neurons in different parts of the circuit. The many sources of non-linearities in neurons may affect visual processing very differently depending on which node of the circuit these non-linearities occur. We have addressed this question using intracellular recordings, agnostic of the definition of RF or the boundary between a center or surround. We have systematically explored the visual space from which synaptic input is triggered in each neuron using stimuli of varying diameters. We found that the majority of supragranular complex cells integrate synaptic input highly sublinearly while infragranular complex cells are very close to linear, while simple cells (regardless of laminar origin) are close to linear.

To explore mechanisms underlying the large difference in linearity of spatial integration between cells, we used a comprehensive large-scale recurrent spiking model of cat V1. Our modeling results suggest that a plausible mechanism for the difference between integration properties of supra- vs. infra-granular layers is differential distribution of long-range horizontal connection patchy terminals between those targeting excitatory vs. inhibitory neurons.

## Materials and Methods

### Subjects

Experiments were performed on 31 adult male cats (2.5-3.5 kg). All experiments were conducted in accordance with the ethical guidelines of the National Institutes of Health and with the approval of the Institutional Animal Care and Use Committee at the University of Pennsylvania.

### Surgical protocol

Surgical methods were as previously reported (Taylor et al., 2018). Animals were sedated in the animal facility with an intramuscular injection of Dexmedetomidine (0.01-0.03 mg/kg) and Buprenorphine (0.02-0.04 mg/kg). Subsequently, animals were brought to the lab and anesthetized with isoflurane (2-3%) and fentanyl i.v. (3-10 ug/kg/hr) for the duration of the experiment (8-10 hrs). Animals were paralyzed with gallamine triethiodide (Flaxedil, 3 mg/kg/hr)) and artificially ventilated, keeping the end-tidal CO2 concentration at 3.9 ± 0.2%. Heart rate, blood pressure, and EEG were monitored throughout the experiment, and the rectal temperature was maintained at 37-38°C with a heating pad. To expose visual cortex, a craniotomy centered at Horsley Clarke coordinates posterior 4.0 and lateral 2.0 was performed and the underlying dura was removed. The stability of recordings was improved by a bilateral pneumothorax, drainage of the cisterna magna, and hip suspension, and by injecting a warm solution of agar (3.5-4%) between the dura and the brain.

### Intracellular recordings

The results described here are based on intracellular recordings from 33 complex cells and 16 simple cells in Area 17. Sharp electrode recordings were performed with glass micropipettes (50-80 MΩ) filled with 3M potassium acetate (KAc). Access resistance of each pipette was compensated online (bridge balancing). All cells had a stable resting membrane potential (Vm) more negative than -60 mV. The average input resistance was 44 ± 15 Mohms (mean ± SD). Only data sets with stable baseline Vm and Rin (<20% change from initial values) throughout the duration of visual stimuli were included. Intracellular recordings were obtained from stereotaxic coordinates AP =-1 to -3 and ML = 0.4 to 0.8.

After a successful penetration, the Vm and input resistance (Rin) became stable within a few minutes and remained so for the duration of the recording (resting Vm: simple cells (n=16) = -68± 6 mV; complex cells (n=33) = -70 ± 8 mV; Rin = 44 ± 15 Mohms; recording time between 30 and 75 mins, n= 49; Values here and below are mean ± SD, unless indicated). We then mapped the cell’s receptive field (RF) by correlating the spikes and Vm with 2D white noise. Visual stimuli were then centered on the RF.

### Visual stimulation

The corneas were protected with neutral contact lenses after dilating the pupil with 1% ophthalmic atropine and retraction of the nictitating membrane with 1% phenylephrine HCl (Neo-Synephrine). Spectacle lenses were chosen by the tapetal reflection technique to obtain a sharp focus of stimuli on the retina. The position of the monitor was adjusted to center the area centralis on the screen. Stimuli were presented on an Image Systems model M09LV monochrome monitor operating at 125 frames/s, a spatial resolution of 1024×786 and mean luminance of 47 cd/m2. The screen subtends 36 by 27° of visual field (28.7 pixels per degree), and lookup tables were linearized for a contrast range of ±100%. Stimuli were generated by custom software writing to the framestore portion of a Cambridge Research Systems visual stimulus generation card mounted in a conventional PC. Computer-assisted hand-plotting routines were used with every cell to provide initial estimates of critical parameters, especially the cell’s RF location and orientation and spatial frequency preferences. Final analysis was performed offline in IGOR (Wavemetrics, OR, USA) from records stored on a Neuralynx system, which included Vm, injected current, and stimulus marks, all sampled at 32 kHz. To map 2D spatial receptive fields, pre-computed frames of low-pass filtered ternary 2D dense noise were presented as 16×16 squares in space covering 2×2° to 4×4°, with a frame duration of 8 ms. Visual stimuli consisted of drifting gratings with optimal spatial frequency and orientation for each cell and at 50% contrast. Gratings were in the shape of 10 to 15 disks of increasing diameter and annuli (“rings”) with a large outer diameter (usually 16 to 20 degrees and an inner diameter equal to that of the disks. Stimuli were presented in random order until all pair diameters (disks and rings) were presented at least 10 times.

### Functional cell classification

Cells were classified as simple or complex based on two criteria, following Cardin et al. (2010). First, we measured the relative modulation of spike trains evoked by an optimized patch of drifting sinusoidal grating. If the response at the fundamental temporal frequency of the stimulus exceeded the average (DC) response, the cell was classified as simple. Otherwise, the cell was classified as complex. Second, we estimated the one- and/or two-dimensional spatiotemporal weighting functions. Cells exhibiting non-overlapping regions excited by bright and dark stimuli were classified as simple. Cells exhibiting excitatory responses (Vm depolarization) to bright and dark stimuli throughout their receptive fields were classified as complex. These two measures yielded the same functional classification.

### Spiking and membrane potential analysis

For Vm measurements, spikes were removed by first determining the time at which spike threshold was reached and then extrapolating the membrane potential values from that point to when the spike repolarized back to the spike threshold level. Following offline spike removal, the Vm was smoothed with a five-point running average. Measurements of the amplitude of the Vm responses were made from the average response to a stimulus across all trials. The mean of the Vm response (VmDC) was measured as the mean of the averaged cyclegram. VmF1 was calculated as the amplitude of the sinusoidal fit to the average Vm cyclegram. We quantified synaptic noise as the standard deviation of the Vm during the single trial cyclegrams (VmSD). Each single trial cyclegram was divided in three equal parts, the VmSD calculated for each and the value from the three averaged to generate that single trial VmSD. The VmSD presented in the results is the average of all trials for each given disk or ring stimulus.

### Suppression index

We quantified the amount of suppression elicited by large disks with a suppression index (SI) by computing the difference between the maximum response *D*_peak_ elicited by a disk stimulus and the response elicited by the largest disk *D*_largest_, and then dividing it by *D*_peak_:

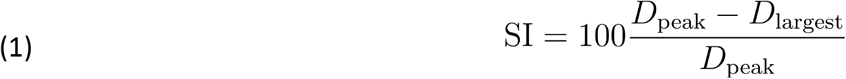

The result is multiplied by 100 so that it quantifies the percentage of suppression. The responses are calculated after subtracting the baseline activity.

### Residual

Residuals were computed as the ratio between the response *R*_peak_ elicited by a ring stimulus with an inner diameter equal to the diameter of the disk which elicited the largest response and *D*_peak_:

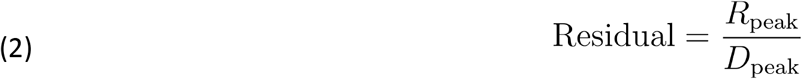

### Non-linearity index

We computed the non-linearity index (NLI) by calculating the difference between *D*_peak_ and the largest component of the algebraic sum of the response to disk and ring pairs with corresponding diameters and inner diameters *D*_*i*_ + *R*_i_ and then dividing it by *D*_peak_:

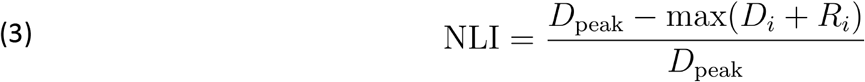

Where *i* represents the index of the disks and rings stimuli.

### Linearity index

The linearity index (LI) is computed for each index *i* of stimuli pair as:

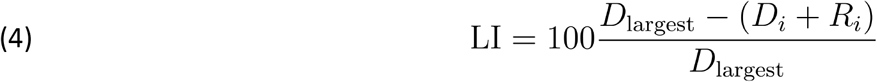

### V1 model

The V1 model presented here is adapted from Antolík et al. (2024). It is composed of two cortical layers, layer 4 and 2/3, which both cover 5 mm^2^ of cortical space and each contain 37 500 excitatory neurons and 9 375 inhibitory neurons (Fig. 8A). Layer 4 receives its feedforward input following a Gabor distribution from a model of the lateral geniculate nucleus (LGN) covering 6° x 6° of visual field and containing 14 400 neurons with center-surround receptive fields. Layer 4 intra-laminar connectivity is local and follows a push-pull scheme. Layer 2/3 inhibitory connectivity is also local whereas excitatory neurons in layer 2-3 send both local and long-range functionally-biased connections. The detailed description of the model has been provided in Antolik et al. (2024). The only difference between the model presented here and its predecessor is in layer 4 inhibitory neurons having higher membrane resistance (327 MΩ vs 300 MΩ in the original model).

### Neuron modeling

Every cortical neuron is modeled as an exponential integrate-and-fire neuron (Gerstner and Brette, 2009). The time course of their membrane potential is modeled by the following equation:

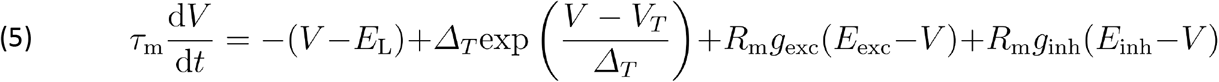

A spike is emitted when the membrane potential of a neuron crosses the threshold value of -40 mV. The neuron then goes through a refractory period of respectively 2 ms and 0.5 ms for excitatory and inhibitory neurons, and its membrane potential is reset to -70 mV. The spike initiation threshold *V*_T_ and the leak potential *E*_L_ are set respectively to -57 mV and -80 mV for excitatory neurons, and to -58 mV and -78 mV for inhibitory neurons. The threshold slope factor *ΔT* of the exponential function is set to 0.8 mV. The excitatory reversal potential *E*_exc_ is set to 0 mV and the inhibitory reversal potential *E*_inh_ is set to -80 mV. For excitatory neurons, the membrane time constant *τ*_m_ and the membrane resistance *R*_m_ are respectively equal to 8 ms and 250 MΩ whereas for inhibitory neurons they are equal to 9 ms and 327 MΩ.

### Input model

The cortical neurons receive input from a model of the LGN where every neuron is represented as a leaky integrate-and-fire model. The LGN neurons receive a current input computed through the convolution of their receptive field modeled as center-surround linear spatio-temporal filter with the stimulus as described in Antolík et al. (2024). An additional gaussian noise current is injected into the LGN neurons so that their baseline spiking response matches experimentally observed values (Papaioannou and White, 1972).

### Thalamo-cortical connections

The input of the LGN neurons to the layer 4 of the model follows a Gabor distribution:

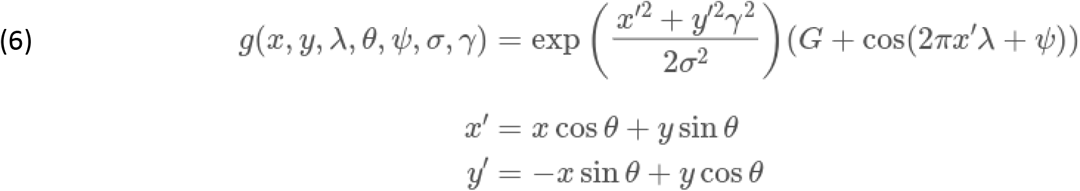

Where both the coordinates *x* and *y*, representing the positions of the neurons, and their spatial phase preference *ψ*, are generated randomly. The size *σ* and the aspect ratio *γ* of the Gabor distribution are set respectively to 0.17° and 2.5. The spatial frequency *λ* of the distribution is equal to 0.8 cycles per degree, whereas its orientation varies for each neuron and depends on its position on a pre-computed orientation map. *G* is set to 0.085 and represents the weight of the Gaussian relative to the Gabor distribution. Layer 4 excitatory cells receive a number of thalamo-cortical afferents drawn uniformly within the range [90,190], whereas this range is [112,168] for inhibitory cells. These intervals lie within the range of anatomical observations (Da Costa and Martin, 2011).

### Cortico-cortical connectivity

Excitatory neurons in layer 4 receive 1000 synapses, whereas those in layer 2/3 receive 2300 synapses. To account for their smaller size, inhibitory neurons receive 40% less synapses. Excitatory and inhibitory synapses form on cortical neurons at a ratio of 4:1. 22% of the excitatory input of layer 2/3 comes from feedforward input from layer 4, whereas 20% of the excitatory input in layer 4 is formed by feedback connections originating from layer 2/3. All the other cortico-cortical synapses are formed by intra-laminar connections. Both cortico-cortical connectivity in layer 4 and inhibitory connectivity in layer 2/3 are local, with the probability density function pdf of synapses forming between pairs of neurons being computed by one distance-dependent component *p*_dist_ and one functional-dependent component *p*_func_:

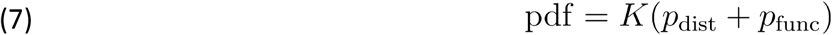

Where K is a normalizing constant defined so that the probabilities sum up to 1. p_dist_ follows a hyperbolic distribution:

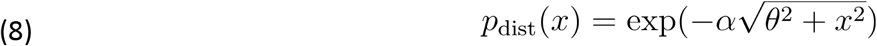

where x is the distance between pairs of neurons, and *α* and *θ* parameters are obtained by fitting the data (see Antolík et al. (2024) for details) from Stepanyants et al. (Stepanyants et al., 2008) and listed in table 1.

**Table 1:** Parameters of the hyperbolic distributions used to generate the local connectivity of the model. Rows refer to pre-synaptic populations, and columns to post-synaptic populations.

| | $\alpha$ | | | | $\theta$ | | | |
| --- | --- | --- | --- | --- | --- | --- | --- | --- |
|  | L4 Exc | L4 Inh | L23 Exc | L23 Inh | L4 Exc | L4 Inh | L23 Exc | L23 Inh |
| L4 Exc | 0.0139 | 0.0148 | 0.0174 | 0.0197 | 207.7 | 191.8 | 154.4 | 131.5 |
| L4 Inh | 0.0126 | 0.0119 |  |  | 237.5 | 256.4 |  |  |
| L23 Inh |  |  | 0.0149 | 0.0150 |  |  | 189.5 | 188.61 |

The recurrent connectivity in layer 4 follows a push-pull scheme: Excitatory connections are more likely to form between neurons with correlated receptive fields (defined from the gabor distribution used for drawing the thalamo-cortical connections), and inhibitory connections are more probable between neurons with anti-correlated receptive fields. Its functional component is then the following:

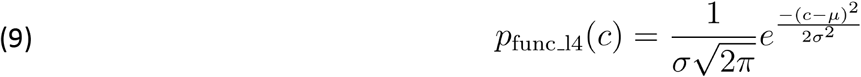

With *c* the level of correlation between the pair of neurons, *σ* = 1.3 radians, and *µ* = 1 when the pre-synaptic neuron is excitatory or -1 when it is inhibitory.

There is no functional component in the inhibitory connectivity in layer 2/3 and *p*_func_ is thus equal to 0 for these connections. The feedforward connectivity to layer 2/3 has a bias for connecting neurons sharing similar orientation preference. Its functional component is:

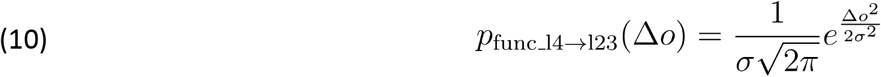

Where Δ*o* is the difference in orientation preference between the pair of neurons based on the pre-computed orientation map of the model, and *σ* is equal to 1.3 radians.

The distance-dependent excitatory connectivity in layer 2/3 has both a local component, with no explicit functional bias, and a long-range component which is modulated by a functional component representing the orientation bias of the horizontal connectivity in this layer:

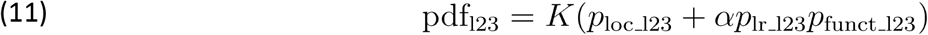

With p_loc_l23_ and p_lr_l23_ corresponding respectively to the local and long-range distance-dependent components, and K a normalizing constant defined so that the probabilities sum up to 1. p_funct_l23_ represents the functional component and follows the same equation as for the layer 2/3 feedforward connectivity. The weight *α* of the long-range component relatively to the local one is equal to 4. Both p_loc_l23_ and p_lr_l23_ follows the following equation:

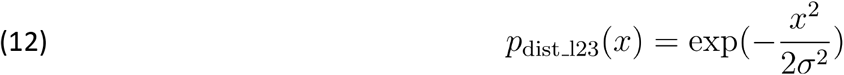

with x the distance between pairs of neurons, and σ equal to 270 mm and 1000 mm respectively for the local and long-range components.

### Synapses

Each synapse is modeled with an exponential decay:

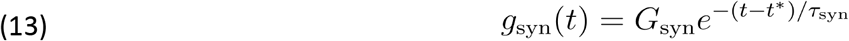

Where G_syn_ represents the weight of the synapses, values for each type of connection is shown in Table 2. t* is the spike time and τ_syn_ is the time constant of the exponential decay, equal to 1.5 ms and 4.2 ms respectively for excitatory and inhibitory synapses.

**Table 2:** Parameters of the model connectivity.

| Source | Target | Incomming synapses per post-synaptic neuron | Weight | Description of the connection probabilities between neurons |
| --- | --- | --- | --- | --- |
| LGN ON | Layer 4 Exc | Uniform within [45,95] interval | 1.2 nS | Gabor distribution |
| LGN OFF | Layer 4 Exc | Uniform within [45,95] interval | 1.2 nS | Gabor distribution |
| LGN ON | Layer 4 Inh | Uniform within [56,84] interval | 1.2 nS | Gabor distribution |
| LGN OFF | Layer 4 Inh | Uniform within [56,84] interval | 1.2 nS | Gabor distribution |
| Layer 4 Exc | Layer 4 Exc | 640 – number of synapses received from the LGN | 0.18 nS | Distance-dependent with ‘Push’ mechanism |
| Layer 4 Exc | Layer 4 Inh | 384 – number of synapses received from the LGN | 0.22 nS | Distance-dependent with ‘Push’ mechanism |
| Layer 4 Inh | Layer 4 Exc | 200 | 1 nS | Distance-dependent with ‘Pull’ mechanism |
| Layer 4 Inh | Layer 4 Inh | 120 | 1 nS | Distance-dependent with ‘Pull’ mechanism |
| Layer 4 Exc | Layer 2/3 Exc | 404 | 1 nS | Distance-dependent with orientation bias |
| Layer 4 Exc | Layer 2/3 Inh | 242 | 1 nS | Distance-dependent with orientation bias |
| Layer 2/3 Exc | Layer 2/3 Exc | 1435 | 0.18 nS | Local distance-dependent component<br>Long-range distance dependent component with orientation bias |
| Layer 2/3 Exc | Layer 2/3 Inh | 861 | 0.35 nS | Local distance-dependent component<br>Long-range distance dependent component with orientation bias |
| Layer 2/3 Inh | Layer 2/3 Exc | 460 | 1 nS | Distance-dependent |
| Layer 2/3 Inh | Layer 2/3 Inh | 276 | 1 nS | Distance-dependent |
| Layer 2/3 Exc | Layer 4 Exc | 160 | 0.18 nS | Distance-dependent with orientation bias |
| Layer 2/3 Exc | Layer 4 Inh | 96 | 0.22 nS | Distance-dependent with orientation bias |

The delays of the cortico-cortical synapses are computed as the sum of a constant and of a distant-dependent component with a propagation delay of 0.33 s/m. The constant component is equal to 1.4 and 0.5 ms for excitatory connections targeting respectively excitatory and inhibitory neurons, and to 1 ms for inhibitory connections. The delays of the thalamic afferents are independent from distance and drawn randomly from an interval of [1.4, 2.4] ms. Every synapse of the model exhibits short-term depression following the Tsodyks-Markram model (Tsodyks et al., 2000), with parameters for different types of connections shown in Table 3.

**Table 3:** Short-term plasticity parameters of the connections of the model.

| Source | Target | Effective use of the synaptic resources ($u$) | Recovery time constant ($\tau_{rec}$) | Synaptic current time constant ($\tau_{psc}$) |
| --- | --- | --- | --- | --- |
| LGN ON | Layer 4 Exc | 0.75 | 125 ms | 1.5 ms |
| LGN OFF | Layer 4 Exc | 0.75 | 125 ms | 1.5 ms |
| LGN ON | Layer 4 Inh | 0.75 | 125 ms | 1.5 ms |
| LGN OFF | Layer 4 Inh | 0.75 | 125 ms | 1.5 ms |
| Layer 4 Exc | Layer 4 Exc | 0.75 | 30 ms | 1.5 ms |
| Layer 4 Exc | Layer 4 Inh | 0.75 | 30 ms | 1.5 ms |
| Layer 4 Inh | Layer 4 Exc | 0.75 | 70 ms | 4.2 ms |
| Layer 4 Inh | Layer 4 Inh | 0.75 | 70 ms | 4.2 ms |
| Layer 4 Exc | Layer 2/3 Exc | 0.75 | 30 ms | 1.5 ms |
| Layer 4 Exc | Layer 2/3 Inh | 0.75 | 30 ms | 1.5 ms |
| Layer 2/3 Exc | Layer 2/3 Exc | 0.75 | 30 ms | 1.5 ms |
| Layer 2/3 Exc | Layer 2/3 Inh | 0.75 | 30 ms | 1.5 ms |
| Layer 2/3 Inh | Layer 2/3 Exc | 0.75 | 30 ms | 4.2 ms |
| Layer 2/3 Inh | Layer 2/3 Inh | 0.75 | 30 ms | 4.2 ms |
| Layer 2/3 Exc | Layer 4 Exc | 0.75 | 30 ms | 1.5 ms |
| Layer 2/3 Exc | Layer 4 Inh | 0.75 | 30 ms | 1.5 ms |

### Model simulations

We recorded the spiking activity of and the membrane potential of layer 4 and layer 2/3 model cells located within 0.4° of the center of the visual field. Cells were stimulated using sinusoidal drifting gratings of optimal orientation confined in disks and rings similarly to the experimental data. We used 10 values of radii for each stimulus and repeated each stimulus 10 times for 1500 ms. The order of stimuli was randomly shuffled for the presentation. We only selected cells which exhibited a mean firing rate higher than 1 spike/s across the 10 repetitions of at least one instance of the disk stimuli.

### Data analysis

We classified model cells as simple or complex based on their responses to 10 repetitions of an optimized patch of grating of 15° of diameter. If their spiking response at the fundamental frequency of the stimulus exceeded their average response, then the cells were classified as simple, otherwise as complex. We excluded from further analysis simple cells for which the mean membrane potential depolarization was larger than its Fourier component at the fundamental frequency of the stimulus. This resulted in 27 simple cells and 36 complex cells analyzed. We computed the standard deviation of the cells Vm on single trials as the mean of the standard deviations computed using a 100 ms sliding window over the course of their Vm. For simple cells, this was done after subtracting their Vm response at the fundamental frequency of the stimulus.

### Modifications of layer 2/3 connectivity parameter

To study the impact of the parameters of long-range connectivity of the layer 2/3 model on the spatial integration properties of cells, we modified the spatial range of the long-range component of recurrent excitatory connections targeting excitatory neurons (σ in eq. (12)), while keeping fixed the corresponding parameters for connections targeting inhibitory neurons. We chose values of 1250 mm (Fig10, Broad E→E range), and 740 mm (Fig10, Narrow E→E range). Similarly, we also varied the level of orientation bias in the long-range connectivity targeting of excitatory neurons (σ in eq. (10)), choosing values of 0.7 radians (Fig10, High E→E or. bias) and 3.5 radians (Fig10, Low E→E or. bias).

## Results

We used intracellular recordings from the primary visual cortex (V1) to measure the membrane potential (Vm) and spike output in response to visual stimuli. Visual stimuli consisted of optimal drifting gratings confined to (1) circular patches (“disks”) of increasing diameter and (2) matched circular annuli (“rings”) with inner diameters identical to those of the corresponding disk and fixed outer diameter (Fig 1B). All stimuli were randomly interleaved.

**Fig. 1.**
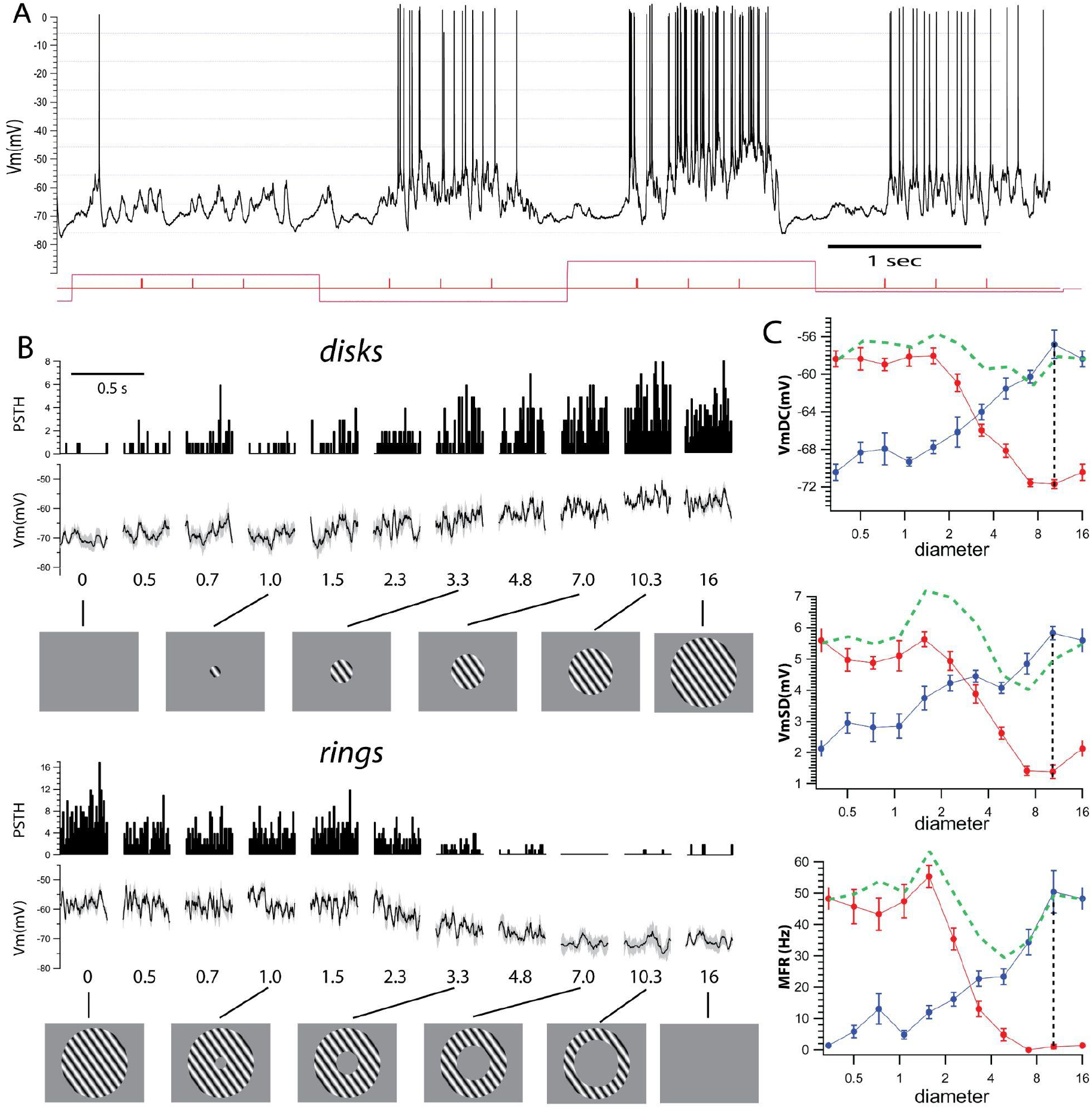
Example infragranular complex cell (IGC) and visual stimulation protocol. **A**. Example intracellular trace from an IGC cell responding to varying visual stimuli. The cell responds with sustained depolarization and firing rate to the drifting gratings. **B**. Cyclegrams and PSTHs show increased average depolarization and spike output with the increase in disk diameter (top two rows, diameter in degrees of visual space is indicated below each cyclegram). Responses to rings (bottom two rows) show a decrease of response towards resting levels with the increase in the inner diameter (indicated below), outer diameter of the rings is 16 degrees. **C**. Quantification of the response shows sustained increase in all three parameters, VmDC, VmSD and MFR for disks (blue) and reduction for rings (red). Vertical dotted lines show that at the peak response to a disk there is no response to a ring with that inner diameter. Green dotted lines show the algebraic sum of disk and ring pairs.

Cell depth was read from the micromanipulator and compared to the position of a few cells filled with neurobiotin. Although the manipulator reading and the histological position of the recovered cells was accurate within 200 μm, the curvature of the posterior gyrus of the cat brain adds uncertainty to the reading. Thus, rather than the exact position of the cells, we classified our complex cells broadly as either supragranular layers 2-3 (L2-3, 200 to 700 μm) or infragranular layers 5-6 (L5-6, 900 to 1600 μm). The distance between them is larger than the error and is separated by consistently finding simple cells between 600 and 900 μm. Following this classification, for population analysis, we will consider three cell categories: complex supragranular (SGC), complex infragranular (IGC) and simple cells from granular (GS) and infragranular (IGS) layers.

### Infragranular complex cells

An example intracellular recording of an IGC cell had a resting Vm around -70 mV and responded with varying levels of sustained depolarization and spike output to three cycles of the drifting grating (Fig. 1A, cycle starts indicated by tick marks). For each stimulus condition, we averaged the Vm and accumulated the spikes in peristimulus spike histograms (PSTH) for the duration of the cycle (cyclegrams, Fig. 1B). The Vm and PSTH cyclegrams showed a progressive depolarization and increase in firing rate, respectively, to a maximum for a disk of 10.3 degrees, followed by a small suppression (Fig. 1B, disks). Corresponding ring stimuli triggered progressively smaller responses reaching resting values at an inner diameter of 7.0 degrees (Fig. 1B, rings).

We quantified the response of complex cells from the cyclegrams by calculating (i) the mean depolarization of the Vm (VmDC), (ii) the mean standard deviation of the Vm (VmSD, measured from single trials that make up each average cyclegram, see Methods), which represents synaptic noise, and (iii) the mean firing rate (MFR) calculated as the number of spikes per single cycle divided by the duration of the cycle (Fig. 1, right column, disks in blue, rings in red). Response quantification (Fig. 1C) showed that the maximum response for a disk of 10.3 degrees consisted of a 13.6 mV VmDC depolarization from rest (rest = -70.4 ± 0.9 mV, peak = -56.8 ± 1.5 mV), an increase in VmSD of 3.7 mV (rest= 2.1 ± 0.3 mV, peak = 5.8 ± 0.2 mV), and an increase in MFR of 49 Hz (rest =1.3 ± 1.0 Hz; peak = 50.4 ± 6.7 Hz).

The ring with the inner diameter equal to the size of the disk of maximum response (10.3 degrees, Fig. 1C red traces, indicated by vertical dotted line) did not trigger a response in either the VmDC (−71.7 ± 0.5 mV), VmSD (1.4 ± 0.2 mV) or MFR (1.0 ± 0.7 Hz), none of these values were different from rest (p > 0.5, Kolmogorov-Smirnov test). Moreover, in this cell, all three measures reached resting levels for a ring with even a smaller inner diameter of 7.0 degrees, suggesting that most of the excitatory input that drives the cell is encompassed by the disk of 10 degrees and that there is no input triggered from stimuli beyond this area.

### Supragranular complex cells

In the example supragranular cell in Fig. 2, Vm cyclegrams showed increasing depolarization from rest with increasing disk diameter up to a peak value (at 1.8 degrees) followed by a small suppression (Fig. 2B, disks, Vm). This behavior was paralleled by the spike output (Fig. 2B, PSTHs). Conversely, the Vm and spike responses to the rings decreased with the increasing inner diameter, until no depolarization or spike output was triggered for rings with inner diameters larger than 3.8 degrees (Fig. 2B, rings) Quantification of the cyclegrams in this cell showed that the VmDC increased steadily with increasing disk diameter to a maximum of 18 mV (disk = 1.8 degrees; rest Vm = -77.3 ± 1.3 mV; peak Vm = -59.0 ± 1.2 mV), followed by small suppression for larger diameters (−61 ± 0.6 mV at the largest disk diameter of 10 degrees). This value of suppression represents a reduction of 2 mV, or 11%, of the peak value. The VmSD increased from 0.7 ± 0.2 mV at rest to a peak of 2.9 ± 0.1 mV (at 1.0 degrees) followed by a small response suppression (2.6 ±0.2 mV at 10 degrees, a reduction of 10% of the peak). The MFR increased to a peak of 10.5 ± 1.4 Hz (rest = 0 Hz, peak = 1.5 degrees) followed by strong suppression (2.3 ± 0.7 Hz at 10 degrees, a reduction of 78% of the peak). The much larger suppression of the spike response reflects the well-known steep relationship between Vm and spikes, by which small changes in Vm lead to large changes in spike rate.

**Fig. 2.**
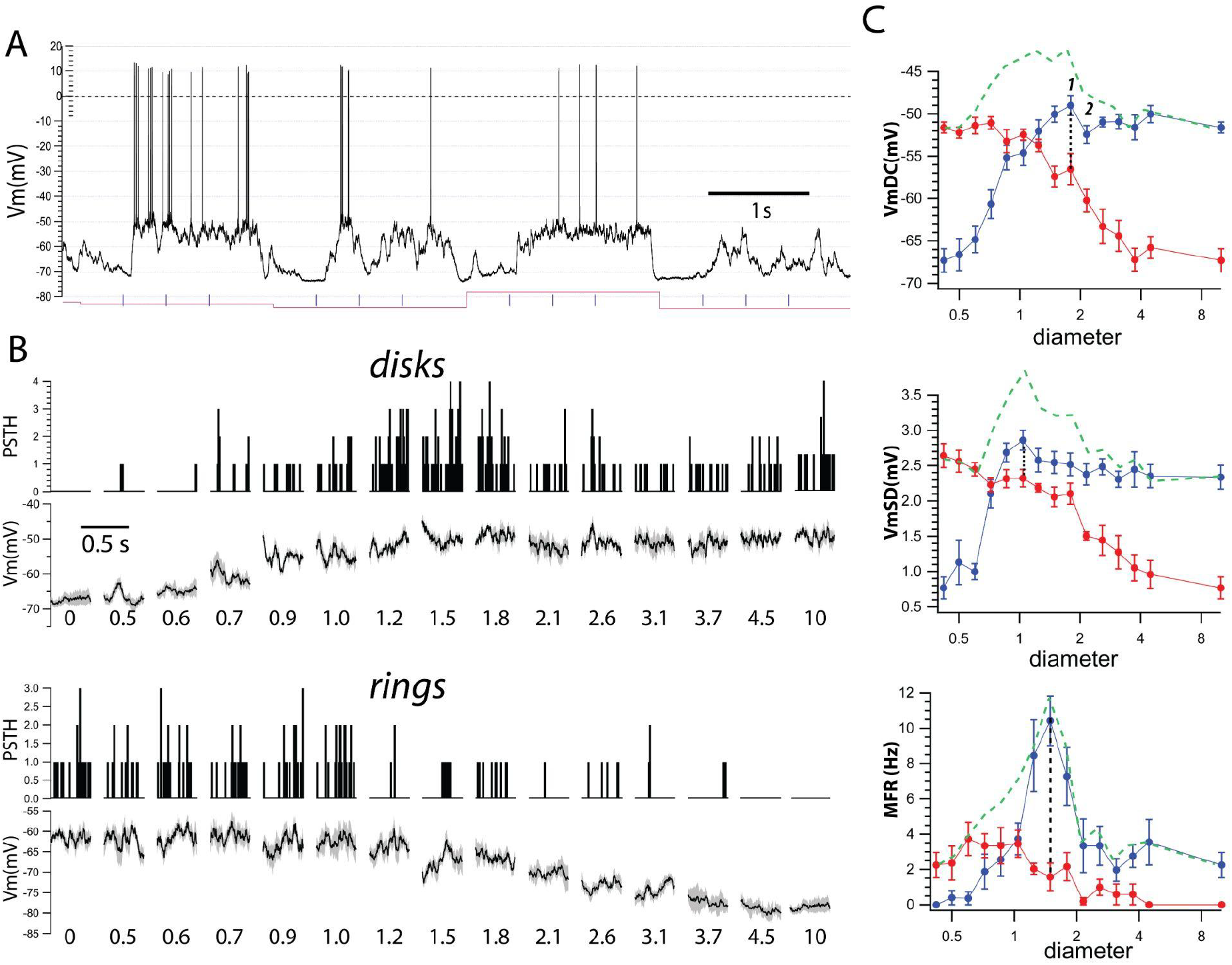
Example supragranular complex cell (SGC). **A**. Intracellular recording from a supragranular complex cell. The example shows a snapshot of cellular activity during visual stimulation. Red line indicates changes in visual stimuli. Tick marks represent the start of drifting grating cycles. The membrane potential (Vm) shows sustained depolarization and spike firing in response to drifting gratings of different diameters. **B**. Averaged Vm around the drifting grating cycles (cyclegrams) show depolarization with increasing diameter of disks (disks, top row, diameter indicated in degrees of visual space) and progressive repolarization to resting level with increasing inner diameter of rings (rings, bottom row). In this and following figures, zero diameter signifies a uniform gray screen for disks, and a large solid disk, i.e., a ring with zero inner diameter. **C**. Quantification of the responses show increased response as a function of disk diameter (blue) and reduction with increasing inner diameter of the rings (red). At the peak disk response (point #1) there was a large response to the corresponding ring (dotted lines). A ring with inner diameter of a disk that causes suppression (point #2) still causes a large response. Green dashed lines are the algebraic sum of the disk and ring pairs.

In stark contrast with the infragranular cell shown in Fig. 1, the ring with an inner diameter equal to that of the disk triggering the peak VmDC (1.8 degrees, outer diameter = 10 degrees, point #1), thus entirely within the suppressive area, caused a large depolarization to -66.5 ± 1.8 mV (9 mV from rest or 50% of the peak response). Even a ring of a larger inner diameter (2.1 degrees, point #2), i.e. further into the suppressive surround (Fig. 1), still elicited a depolarization of 7 mV from rest (40% of the peak depolarization). This was similar for VmSD, where the ring with inner diameter equal to the disk triggering the peak VmSD (see above) caused 2.3 mV of synaptic noise (80% of the peak) and the next larger inner diameter ring (1.2 degrees), still triggered a 2.2 mV VmSD, or 75% of the peak. Thus, even though disks with diameter beyond those eliciting the maximum response cause suppression of both VmDC and VmSD, a ring that exclusively stimulates the suppressive area triggers strong depolarization and large synaptic noise.

### Simple cells

The example simple cell from layer 4 (Fig. 3) showed a characteristic sinusoidal modulation of the Vm and spikes at the frequency of the drifting gratings. Vm cyclegrams and PSTHs showed that the amplitude of the response modulation increased as a function of disk diameter up to a peak at 1.9 degrees (Fig. 3B, disks). The corresponding rings showed a decrease in modulation of Vm and PSTH as a function of inner ring diameter, reaching resting values for the diameter equal to the disk peak response (Fig. 3B, rings). Thus, a ring with inner diameter equal to the disk that triggered maximum response modulation caused no response.

**Fig. 3.**
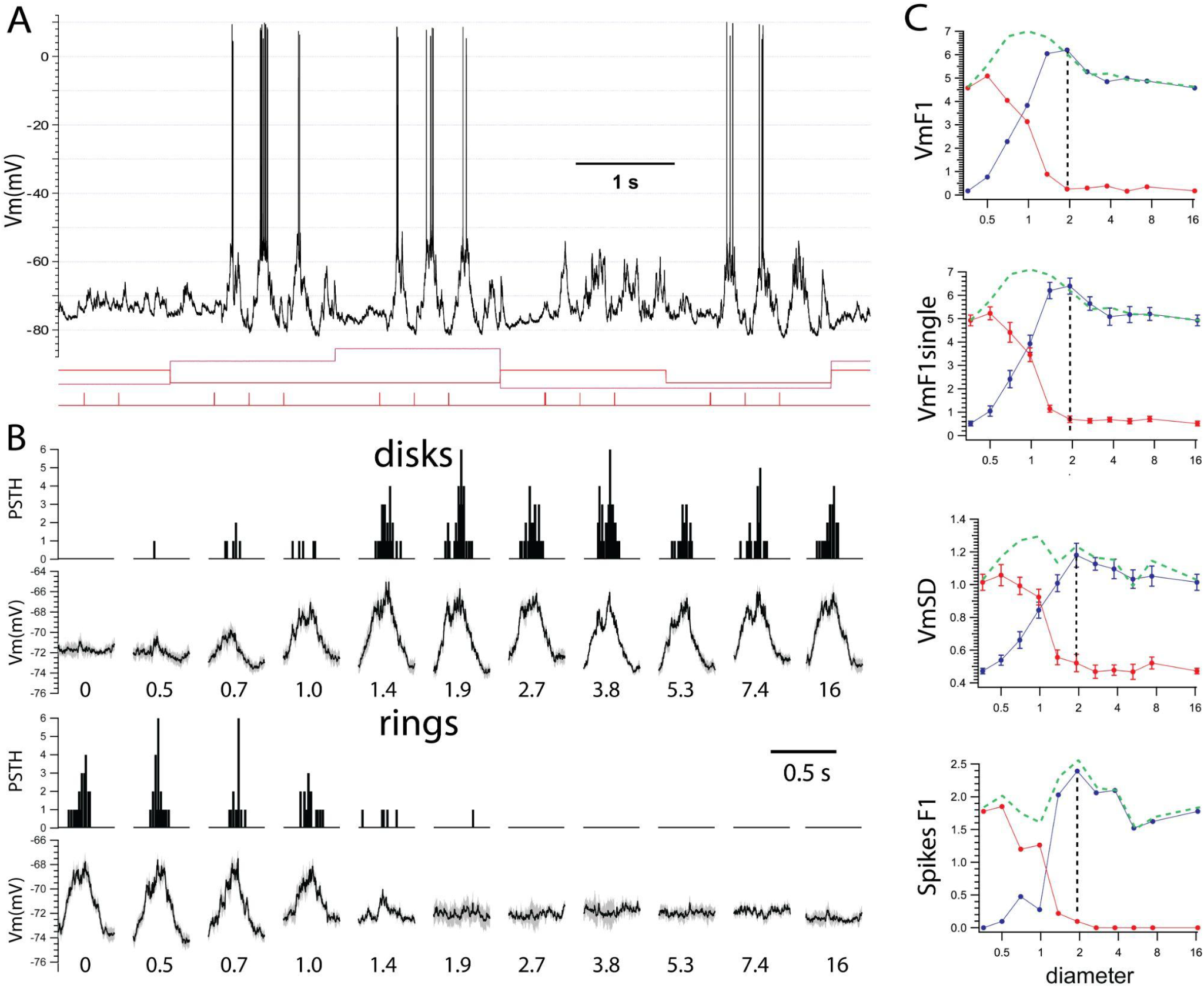
Example simple cell. **A**. Intracellular recording from a simple cell displaying a characteristic temporal modulation of Vm and spikes to drifting gratings. Varying stimulus conditions indicated by changing levels of red line. **B**. Cyclegrams show the phase dependent modulated responses that increase in amplitude with disk diameter (top two rows) and correspondingly decrease with increasing inner diameter. **C**. Quantification of the cyclegrams show that a ring (red) with an inner diameter equal to that of the disk (blue) triggering maximum response (dotted line), has no response. Green dotted lines show the algebraic sum of disk and ring responses. Values discussed in text.

We quantified the response of simple cells as (i) the amplitude of a sine wave at the temporal frequency of the drifting grating fit to the average response Vm (VmF1), (ii) the mean standard deviation of the Vm (VmSD) around the sine-modulated Vm, calculated from single trials. Spike output was quantified as the amplitude of a sine wave fit to the PSTH (spikes F1).

In the simple cell in Fig. 3, the VmF1 increased with the diameter of the disk by 6.2 mV (Fig. 3C, blue, rest F1 = 0.6 mV, peak = 6.8 mV, disk = 1.9 degrees) followed by a suppression of 19% of the peak (F1 = 5.5 mV at 16.5 degrees). Similarly, the VmF1 calculated from single trials (VmF1single) reached a peak of 6.7 ± 0.3 mV for a disk of the same diameter of 1.9 degrees. In response to rings (in red) of increasing inner diameter, VmF1 decreased rapidly and reached a baseline for an inner diameter also of 1.9 degrees. The VmSD and spikes F1 behaved similarly, and both reached a maximum of 1.2 ± 0.1 mV (rest = 0.46 mV) and 2.4 Hz (rest = 0 Hz), respectively, in response to the 1.9 degree disk and returned to baseline levels for rings of the same inner diameter.

Thus, simple cells show no remaining excitatory drive beyond the region eliciting maximum response similarly to infragranular cells.Simple cells.

The example simple cell from layer showed a characteristic sinusoidal modulation of the Vm and spikes at the frequency of the drifting gratings (Fig. 3). Vm cyclegrams and PSTHs showed that the amplitude of the response modulation increased as a function of disk diameter up to a peak at 1.9 degrees (Fig. 3B, disks). The corresponding rings showed a decrease in modulation of Vm and PSTH as a function of inner ring diameter, reaching resting values for the diameter equal to the disk peak response (Fig. 3B, rings). Thus, a ring with inner diameter equal to the disk that triggered maximum response modulation caused no response.

We quantified the response of simple cells as (i) the amplitude of a sine wave at the temporal frequency of the drifting grating fit to the average response Vm (VmF1), (ii) the mean standard deviation of the Vm (VmSD) around the sine-modulated Vm, calculated from single trials. Spike output was quantified as the amplitude of a sine wave fit to the PSTH (spikes F1).

In the simple cell in Fig. 3, the VmF1 increased with the diameter of the disk by 6.2 mV (Fig. 3C, blue, rest F1 = 0.6 mV, peak = 6.8 mV, disk = 1.9 degrees) followed by a suppression of 19% of the peak (F1 = 5.5 mV at 16.5 degrees). Similarly, the VmF1 calculated from single trials (VmF1single) reached a peak of 6.7 ± 0.3 mV for a disk of the same diameter of 1.9 degrees. In response to rings (in red) of increasing inner diameter, VmF1 decreased rapidly and reached a baseline for an inner diameter also of 1.9 degrees. The VmSD and spikes F1 behaved similarly, and both reached a maximum of 1.2 ± 0.1 mV (rest = 0.46 mV) and 2.4 Hz (rest = 0 Hz), respectively, in response to the 1.9 degree disk and returned to baseline levels for rings of the same inner diameter.

Thus, simple cells show no remaining excitatory drive beyond the region eliciting maximum response similarly to infragranular cells.

### Measures of spatial integration: 1. Suppression

We observed that beyond the maximum response, increasing the diameter of the disk caused some level of suppression. We quantified it with a suppression index (SI, see methods).

Our values of synaptic response suppression (Fig. 4A, Medians of SI are: VmDC=19.4%; VmF1=21.6%; VmSD complex cells = 17.3%; VmSD simple cells = 8.9%; MFR = 49.7%; Spikes F1 = 39.5%) were always above zero, indicating that in all cells there was some degree of suppression. SI values for spikes were larger than for Vm as expected for the steep relationship between Vm and spikes (Carandini and Ferster, 2000; Wilent and Contreras, 2005; Cardin et al., 2010). The values of SI show that in all cells the increase in disk diameter beyond the maximum response brought about a mechanism of suppression that reduced the response.

**Fig. 4.**
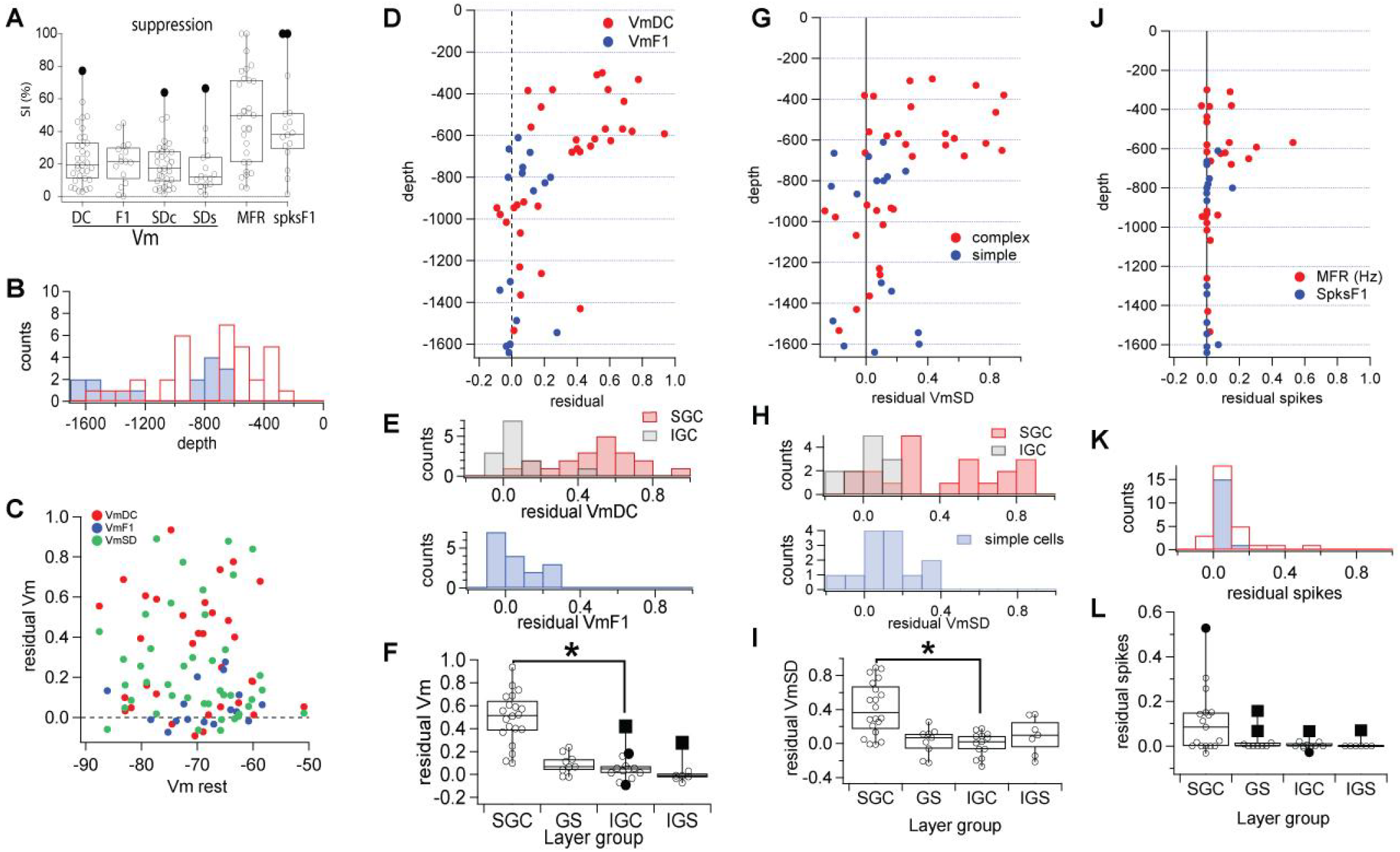
Residuals of Vm and spikes. **A**. Suppression index (SI) of complex and simple cells. All cells showed some degree of suppression of the Vm response up to 60%. All cells showed suppression of the spike response up to 90%. No differences between supra or infragranular or complex and simple cells.**B**. Number of cells recorded as a function of depth. **C**. Residuals versus resting Vm show no relationship and, therefore, not supporting a strong role for GABAergic inhibition in the suppression mechanism. **D**. The residual Vm for all complex (red, VmDC) and simple (blue, VmF1) cells plotted by the depth of the cell. **E**. top panel: Distribution of residual values for complex cells separated by supra (red) and infra (gray) granular location, and for simple cells (blue). **F**. Box and whisker plots show the significant difference of residual VmDC between SGC and IGC cells but not of residual VmF1 among simple cells from granular or infragranular layers.**G**. Residual VmSD for complex (red) and simple (blue) cells plotted according to cortical depth. **H**. Distribution of the residual VmSD for complex cells was bimodal with SGCs (red) showing values above 0.4 while all IGCs (gray) were below 0.4. Simple cells (blue) showed a unimodal distribution with values below 0.4. **I**. Box and whisker plots show that residual VmSD was different between SGC and IGC but not between infra and granular simple cells. **J**. Residual spike output for complex (MFR, red) and simple (SpikesF1, blue) plotted against cell depth. **K**. Value distributions were unimodal. **K**. Box and whisker plots of residual spikes. These were not statistically different.

### Measures of spatial integration: 2. Residuals

Having observed differences in spatial integration between supra- and infra-granular cells, we wished to quantify this effect at the population level. Our population of intracellular recordings (n=49, 16 simple, 33 complex, out of over 70 recordings) was distributed throughout the depth of the primary visual cortex, as read from the micromanipulator, between 300 and 1640 µm (Fig. 4B).

To quantify the amplitude of the input triggered from beyond the area of maximum response, i.e., from the suppressive area, we calculated a unitless normalized ratio which we called “residual” (see Methods). From each cell, residual is the response to the ring with inner diameter equal to the disk that triggered the peak response divided by the peak response, after subtracting the resting level (response to disk of zero diameter). Thus, a residual value of zero indicates no response to the ring while a value of 1 indicates a response to the ring of the same amplitude as the peak response. Values below zero indicate suppression by the ring below rest, while values above 1 signify a response to the ring larger than the maximum, which we never observed.

### 2a. VmDC and F1

The population data showed that the majority of SGC cells had a residual value for VmDC above 0.4 (0.5 ± 0.2, mean ± SD; n =16 of 20 Fig. 4D, red). This is similar to our example cell in Fig. 2 where the residual was 0.5. In contrast, IGC cells had residual VmDC below 0.2 (0.06 ± 0.1, n = 12 of 13, Fig. 4D, red). The only exception was a fast spiking neuron (presumably an interneuron) at 1430 µm depth with a residual VmDC of 0.41 mV. This difference between SGC and IGCcells created a bimodal distribution for the value of residual VmDC (Fig. 4E, red = supragranular, gray = infragranular), which was statistically significant (Fig. 4F, SGC: median = 0.5 mV, interquartile range = 0.3 mV and IGC: median = 0.05 mV, range = 0.06 mV, p < 0.0001, Kolmogorov-Smirnov; p < 0.0001, Mann-Whitney). Simple cells were recorded in middle cortical layers and in infragranular layers, however, regardless of depth, all simple cells showed a residual VmF1 below 0.25 (0.06 ± 0.1, Fig. 4D, blue), generating a unimodal distribution around zero (Fig. 4E, blue). The residual VmF1 of granular and infragranular simple cells was not different (Fig. 4F; GS: median = 0.07, interquartile range = 0.09 and IGS: median = 0.01, interquartile range = 0.04, Kolmogorov-Smirnov and Mann-Whitney p>0.5).

### 2b. VmSD

We measured the standard deviation of the Vm (VmSD) as a proxy for synaptic noise generated by the total synaptic input to the cell. We calculated the residual for the VmSD and plotted its value as a function of cortical depth (Fig. 4G, red). Similar to VmDC, SGC cells showed a wider range of VmSD residual values, with 10 out of 20 above 0.4 (0.4 ± 0.3), while all IGC cells (n=13) had residual synaptic noise smaller than 0.2 (0.0 ± 0.04; Fig. 4G, red). The residual VmSD distribution for complex cells showed a range between -0.3 to 0.9 (Fig. 4H, top histogram), and SGC (red) and IGC (gray) cells were statistically different (Fig. 4I, SGC: median = 0.4 mV, interquartile range = 0.5 mV; IGC: median = 0.0 mV, range = 0.15 mV; p < 0.0001 Kolmogorov-Smirnov; p < 0.0001 Mann-Whitney test). Thus there was significantly higher synaptic input into SGC versus IGC cells triggered from visual stimuli beyond the diameter of maximum response. The residual VmSD of simple cells was distributed between -0.3 and 0.4 (Fig. 4H, blue) and was not different between granular and infragranular layers (Fig. 4I, p>0.5, Kolmogorov-Smirnov and Mann-Whitney).

Thus, our data show that in IGC cells, the stimulus that causes the peak response engages all the excitatory synaptic input to the cell, with little or none originating from visual space beyond the peak, at least as detectable with our intracellular recordings. This is in contrast with SGC cells where the inner diameter equal to the disk that triggers maximum response still causes a strong depolarization and synaptic noise.

### 2c. Spikes

The differences in residual VmDC and VmSD between supra and IGC cells did not, however, translate to significant differences in spike output. In SGC cells, only 9 out of 20 had residual values of mean firing rate (MFR) above zero (0.1 to 0.5; mean = 0.11 ± 0.14, Fig. 4J, red), while IGC cells showed residual MFR below 0.1 (Fig. 4J, red). This difference did not generate a bimodal distribution (Fig. 4K), nor was it statistically significant (Fig. 4L; SGC: median = 0.12 Hz, interquartile range = 0.28 Hz; IGC: median = 0 Hz, interquartile range = 0.021 Hz; p=0.04, Kolmogorov-Smirnov; p=0.5, Mann-Whitney). Simple cells had residual spikes F1 of zero with the exception of 2 granular and one infragranular simple cells (residual < 0.2).

### 2.d. Residual and Vm

One major tenet in vision research is the concept of surround inhibition. This phenomenon may presumably underlie the presence of suppression in all our intracellular recordings past the diameter of maximum response. Among the many circuit mechanisms to implement surround inhibition, one possibility is the direct GABAergic inhibition of the postsynaptic cell being recorded. One hint at the possible presence of inhibition associated with the ring corresponding with maximum response is the presence of negative residuals. Negative residuals indicate that the response is more hyperpolarized than resting membrane potential. However, the synaptic response to GABAergic inhibitory input is Vm dependent, if strong inhibition is part of the suppression triggered by inputs beyond the maximum response, they should correlate with the resting Vm. We plotted our residual values versus resting Vm and found no relationship (Fig. 4C). Furthermore, a key feature of GABAergic inhibition is the large decrease in input resistance which leads to a large reduction in synaptic potential amplitude. Thus, strong inhibition causes a large decrease in VmSD. However, we found no relationship between VmDC or VmSD with the resting Vm (Fig. 4C). Thus, together with the very small values of negative residuals our data strongly disagrees with the presence of strong inhibition from the surround when stimulated alone, at least as recorded from the soma.

Thus, for both VmDC and VmSD we found a population of complex cells in supragranular layers that shows a large depolarizing synaptic input when stimulated exclusively from the area that causes suppression. In addition, cells with no response from the suppressive region did not show inhibition, i.e., negative residual values were above -0.2 and there was no correlation between the residual values of VmDC and VmSD and the resting Vm.

### Measures of spatial integration: 3. Non-Linearity index

The residual values discussed above, assess spatial integration based on responses to a single pair of disk/ring stimuli with matching outer/inner diameter. To quantify the linearity of integration across all stimulus diameters, we defined a non-linearity index (NLI) calculated by subtracting the maximum disk response from the largest algebraic sum of disk and ring pairs and then dividing by the maximum disk response (see Methods).

On the other hand, an NLI above zero indicates that at least one of the algebraic pairwise sums is larger than the measured maximum disk response indicating that, at least for that value the increase in disk response is not paralleled by a decrease for the ring. Therefore, the summation over space is not linear. Finally, a negative NLI means that for all the algebraic sums of disk and ring pair response were smaller than the disk maximum response, indicating that the ring is suppressive, reducing the response of the neuron below rest.

All NLIs were above zero (Fig. 5). The value of NLI for VmDC plotted against cortical depth (Fig. 5A, red), revealed that the majority (n=16/20) of supragranular complex cells (SGC) had values above 0.5, while for 4 cells it was between 0.2 and 0.4. In contrast, infragranular complex cells (IGC) showed NLI values below 0.5 (n=13; Fig. 5A, red). This difference led to a bimodal distribution (Fig. 5B, top panel, SGC = 0.6 ± 0.2, mean ± SD; IGC = 0.3 ± 0.1). The difference in the NLI value for VmDC between SGC and IGC cells was significant (Fig. 5C red; SGC: median = 0.6, interquartile range = 0.16; IGC: median = 0.3, range = 0.2; p < 0.001, Kolmogorov Smirnov; p < 0.001 Mann-Whitney). Simple cells (Fig. 5A, blue) had NLI of the VmF1 between zero and 0.6 (0.4 ± 0.2; Fig. 5B, blue), and showed no difference between granular and infragranular locations (Fig. 5C, p>0.05). Finally, the NLI of SGC VmDC and the NLI of VmF1 of simple cells (from all layers) was statistically different (p<0.001, both KS and Mann-Whitney). Overall the NLI analysis confirms that the SGC cells perform substantially more non-linear spatial summation than IGC cells and that simple cells. We calculated the NLI for VmSD (SGC = 0.6 ± 0.2; IGC = 0.4 ± 0.3, Simple = 0.5 ± 0.3, mean ± SD), and found no differences among cells (simple or complex) or between different layers. Thus, while the residual VmSD does separate complex cells in two categories, the NLI for VmSD does not.

**Fig. 5.**
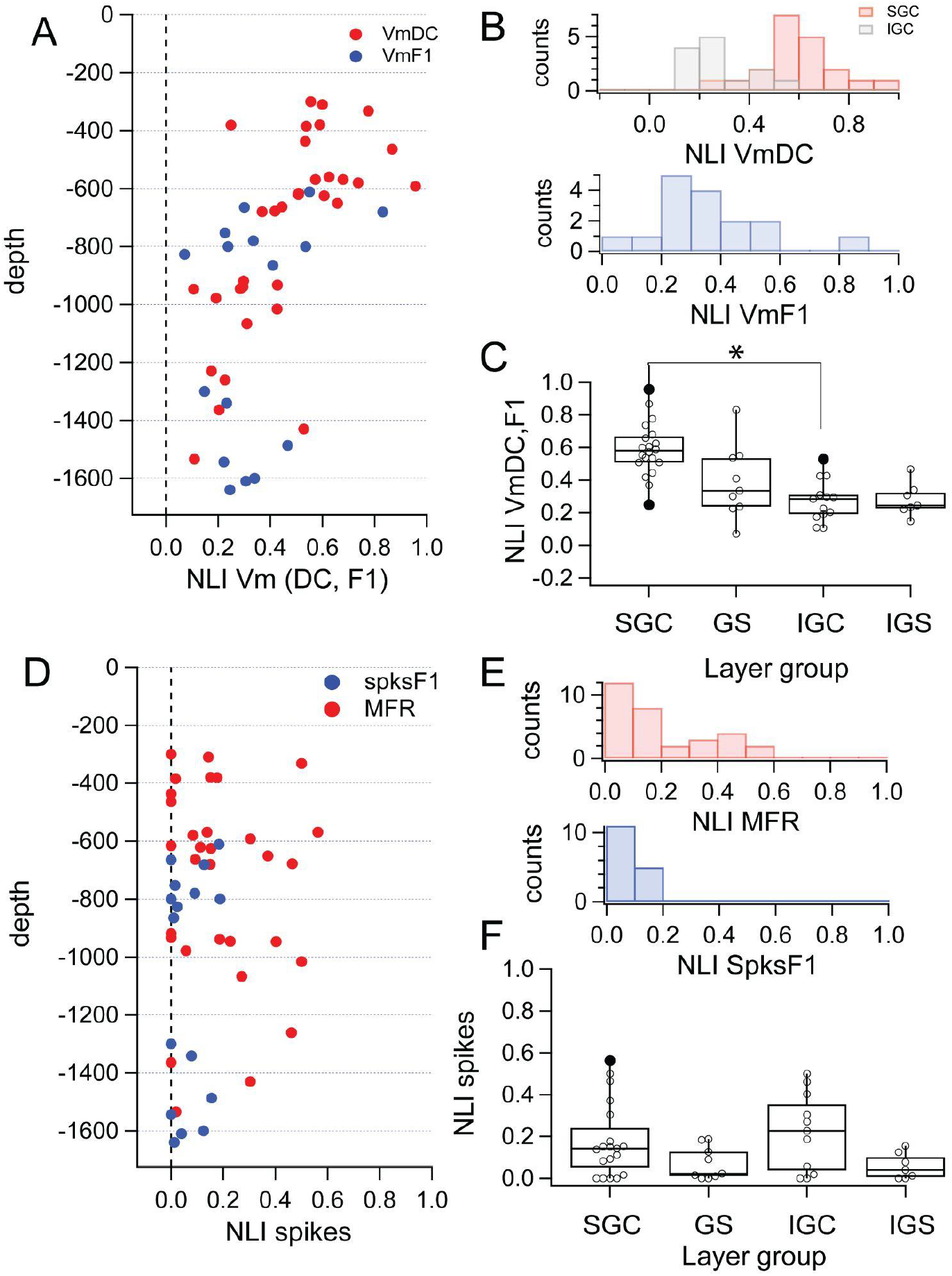
**A**. Value of non-linearity index (NLI) for complex (red) and simple (blue) cells across cortical depth. **B**. Distribution of NLI for SG (red) and IG (gray) complex cells was bimodal while the distribution for simple cells (blue) was unimodal. **C**. SGC and IGC cells were significantly different, but not GS and IGS. **D**. NLI for spikes across the depth. **E**. Distributions of spike output were unimodal for both cell types and cortical depths. **F**. Box and whisker plots show that cell types and cortical depths were not significantly different.

The quantification of the NLI for the MFR of complex cells (Fig. 5D, red) showed many neurons with values spanning from zero to 0.7 (0.2 ± 0.2, mean ± SD ; Fig. 5E, red), but no difference between SGC and IGC neurons (Fig. 5F, red). The spikes F1 of simple cells (Fig. 5D, blue) showed a narrow range between 0 and 0.1 (0.02 ± 0.2; Fig. 5E blue), confirming the result obtained with the residual of spikes F1 response. Furthermore, simple cells in granular and infragranular cells were not different (Fig. 5F).

### Measures of spatial integration: 4. Linearity index

We now address the issue of synaptic input summation in our population of V1 neurons by directly comparing the responses to single stimuli and their algebraic summation with the response to their joint presentation. By our experimental design, each disk and ring stimulus pair sums up to the largest disk presented (see Fig. 1). How does the algebraic sum of the individual responses in each pair (prediction) compare to the response obtained when they are presented together? If the response to the largest disk is smaller than predicted, the cell integrates sublinearly over space, if the response is larger than predicted then the cell integrates input supralinearly. We calculated a linearity index (LI) for each stimulus pair as the difference of obtained (response to largest disk) minus predicted (sum of disk and ring individual responses) responses divided by the response to the largest disk, and multiplied by 100 (see Methods). An LI value of 0% signifies no deviation from linearity (Fig. 6A). Positive and negative LI values indicate supra or sublinear summation, respectively.

**Fig. 6.**
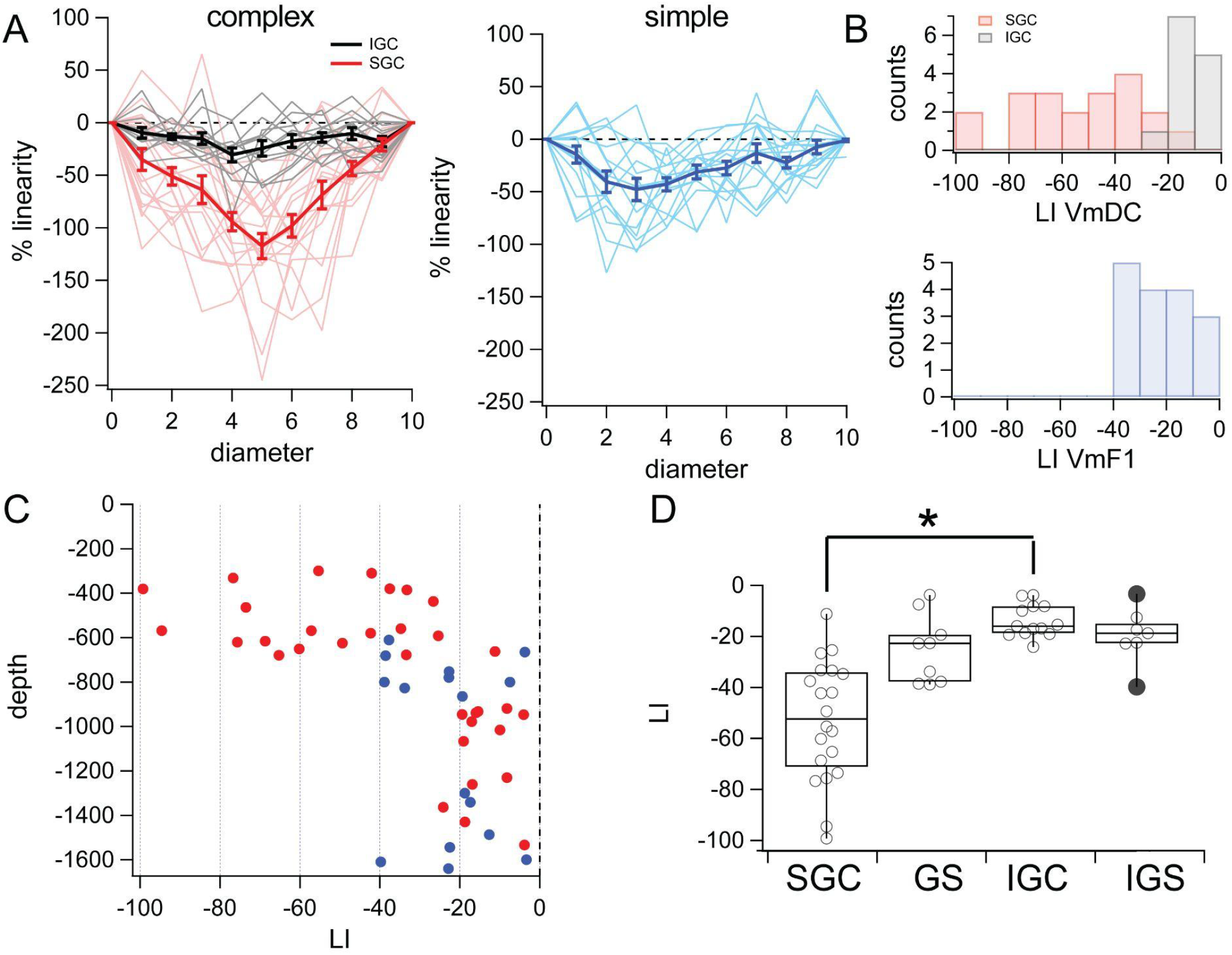
**A**. Thin lines show the LI for each disk-ring pair and each cell for complex (left) and simple (right) cells for all diameters. **B**.The distribution of mean LI for complex cells was bimodal mainly distinguishing supra from infragranular cells, while simple cells show a unimodal distribution. **C**. mean LI of complex and simple cells as a function of cortical depth. **D**. mean LI was significantly different between supra and infragranular complex cells.

The vast majority of values across diameters for both simple and complex cells, were sublinear, i.e., between 0 and -100% (Fig. 6A, thin lines represent individual cells). Averaging the single cells together, showed that SGC cells (Fig. 6A, red) are more sublinear than IGC cells (Fig. 6A, gray), which are very close to perfect linearity. Simple cell values were in between supra and infragranular complex cells (Fig. 6A, blue). In all three classes the average was V-shaped, showing that sublinearity is larger when the individual responses to disk and ring become comparable and the smallest sublinearity occurs when one of the two is very small.

To compare the populations, we calculated the mean of LI across stimuli sizes for each cell. The mean LI values plotted as a function of depth revealed a population of supragranular cells with large deviations from linearity (>40%, Fig. 6B), while all simple cells and IGC were below -40%. The distributions of LI VmDC of SGC and IGC showed minimal overlap (Fig. 6B, top, SGC (red), -53.1 ± 5.3%, mean ± SEM ; IGC (gray) mean= -13.9 ± 6.4%) and was statistically different between the two populations (Fig 6D; SGC: median= -52.4%, interquartile range = 37.0%; IGC: median = -16.0%, IQR = 10.5; p < 0.001, Kolmogorov-Smirnov). The distribution of LI VmF1 for simple cells showed that all values were below 40 % (Fig. 6B, blue, -22.6 ± 3.0%, mean ± SEM) and there was no difference between granular and infragranular simple cells (Fig. 6D, median = -22.6 %, IQR = 20.8%,p>0.05). LI of the VmDC was significantly different between supra and infragranular cells, but the difference in LI VmF1 was not significant between granular or infragranular simple cells. Finally, the LI of SGC VmDC and the LI of VmF1 of simple cells (from all layers) was statistically different (p<0.001, both KS and Mann-Whitney).

### Computational model of non-linear spatial integration

To investigate circuit mechanisms underlying the laminar differences in spatial input integration, we simulated the grating disk and ring protocol in the V1 model from Antolik et al. (2024) (see Methods). The model covers 5 × 5 mm of cortical space and is composed of a layer 4 which receives input from the LGN, and a layer 2/3 which receives feedforward input from layer 4 (Fig. 7A). The connectivity in layer 4 is local and follows a push-pull scheme. The inhibitory connectivity in layer 2/3 is local, but unlike in layer 4 the layer 2/3 excitatory neurons also send long-range horizontal connections moderately biased for iso-oriented targets. The spatial extent and the orientation bias of the long-range connectivity are the same for lateral connections targeting excitatory and inhibitory cells in layer 2/3 (Fig. 7B). Previously, the model was tested against a broad range of stimulation protocols (Antolík et al., 2024) and reproduced both realistic spontaneous activity, contrast-invariant orientation tuning, orientation tuning of subthreshold signals, size tuning, and stimulus-dependent response precision. It also exhibits a realistic laminar distribution of simple and complex cells, which predominate in layer 4 and layer 2/3, respectively.

**Fig. 7:**
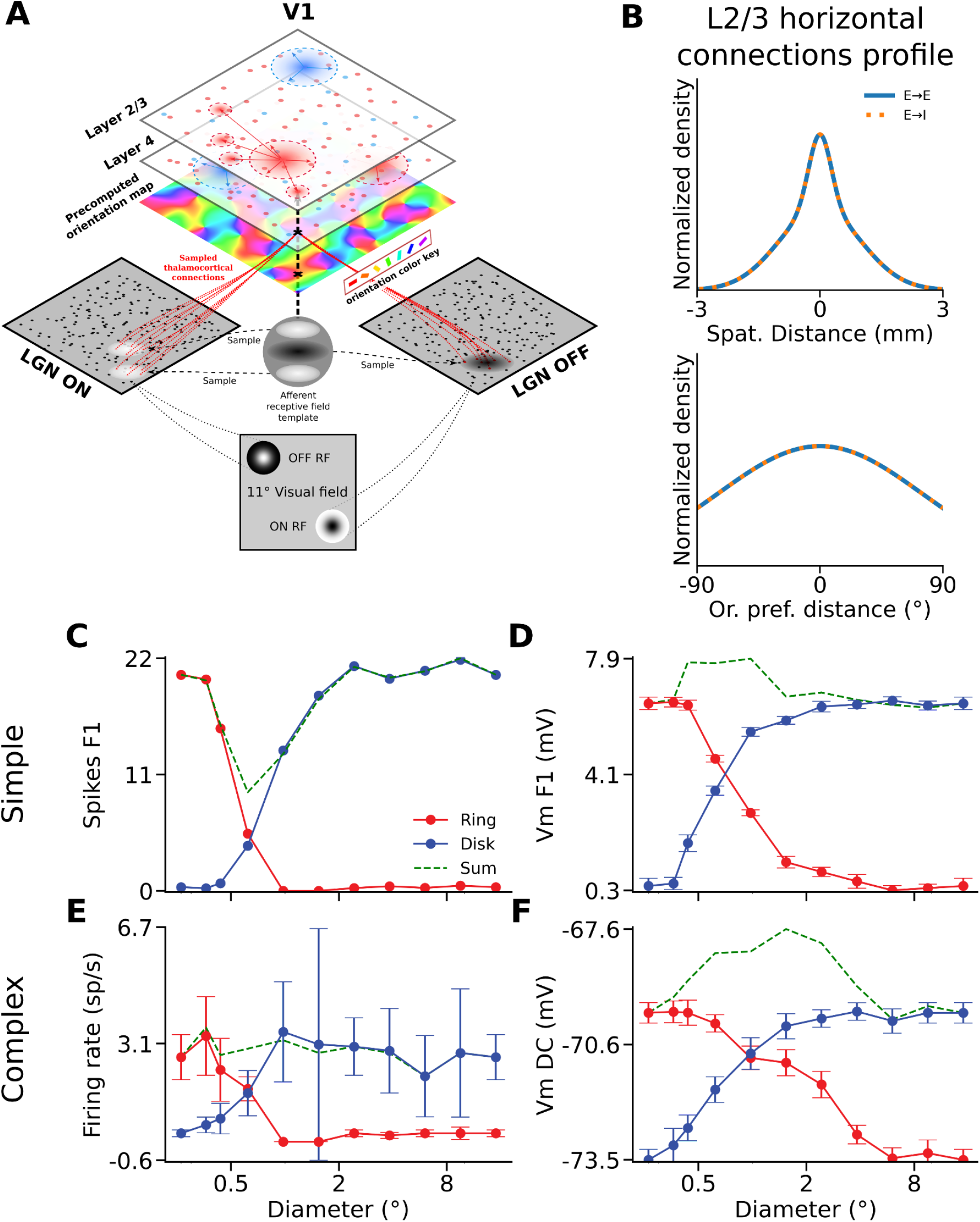
Response of the cells in the model. **A**. Schematic representing the architecture of the V1 model. Thalamic afferents from ON and OFF LGN cells are distributed across post-synaptic layer 4 neurons based on a precomputed orientation map. Local distance-dependent excitatory and inhibitory connections are implemented in each cortical layer. Layer 2/3 receives feedforward input from layer 4 and contains long-range excitatory connections biased for orientation. **B**. Probability distribution of the recurrent excitatory connectivity based on spatial distance (top) and probability distribution of the long-range excitatory connectivity based on the difference in orientation preference (bottom) between pairs of neurons in layer 2/3. Solid blue: excitatory-to-excitatory connections, dashed orange: excitatory-to-inhibitory connections. **C**. Spikes F1 of a model simple cell in response to stimuli of disks varying in diameters (blue) or rings varying in inner diameters (red). The green dashed line represents the sum of the response to both stimuli in respect to baseline.. **D**. Same as **C**, but for its VmF1 response. **E**. Same as **C**, but for the mean firing rate of a model complex cell. **F**. Same as **E**, but for its VmDC response.

We will first demonstrate that the model captures well the linear integration properties of the simple cells vs. nonlinear properties of the supra-granular complex cells revealed in the experiments. We will subsequently study which circuit properties in layer 2/3 control the linearity of the spatial integration, offering an explanation for how neurons in infra-granular layers, despite the presence of long-range lateral connections, exhibit linear spatial integration.

We simulated the disks and rings experiment and quantified spiking and Vm responses of the model neurons as in the in-vivo experiments. Consistent with cat data, we find that the spike output of simple (spikes F1) and complex cells (MFR) dropped to baseline for ring stimuli with inner diameters corresponding to the diameters of the disks that elicited maximum responses (Fig. 7CE). In the example model simple cell (Fig. 7D), the VmF1 saturates for disk diameters at 2.4°, whereas the response to rings with corresponding inner diameter is close to baseline (0.5 mV over baseline, 8 % of the peak). In contrast, the complex cell VmDC (Fig. 7F) saturates for disks with diameters larger than 1.5°, but the cell still exhibits depolarization of 2.5 mV (66% of the peak at 1.5°) for rings with that inner diameter. One important deviation of our model from the cat data is that unlike for spikes, we observe only minor suppression of Vm in most model neurons in response to large disks (Fig 7DF, Complex cells: SI MFR = 25.0 ± 0.7 %, SI VmDC = 1.3 ± 1.3 %, Simple cells: SI Spikes F1 = 9.9 ± 1.4 %, SI VmF1 = 0.7 ± 0.4 %).

To quantify the integration properties of the model neurons at population level, we next calculated the NLIs for all model cells. Both simple and complex cells had, on average, low NLI values for their spikes F1 and MFR respectively (Spikes F1 model simple cells: 0.02 ± 0.05, MFR model complex cells: -0.01 ± 0.04), similarly to the experimental data (Fig. 8AE). The mean NLI for VmF1 and VmDC are in excellent agreement with their experimental counterparts (VmF1 model simple cells: 0.33 ± 0.07; VmDC Model complex cells: 0.52 ± 0.06) (Fig. 8BF), and statistically different between the two cell groups (Simple cells median: 0.32, Complex cells median: 0.50, p < 0.0001, U=23, Mann-Whitney) as in the cat data (Cat simple cells median: 0.30, cat SGC median: 0.59, p = 0.0002, U=41, Mann-Whitney). Overall we note that the range of NLI values is greater in the data than in the model, which is likely due to the lack of neuron-to-neuron variability in some model parameters such as membrane resistance, or number of synaptic inputs from individual projections.

**Fig. 8:**
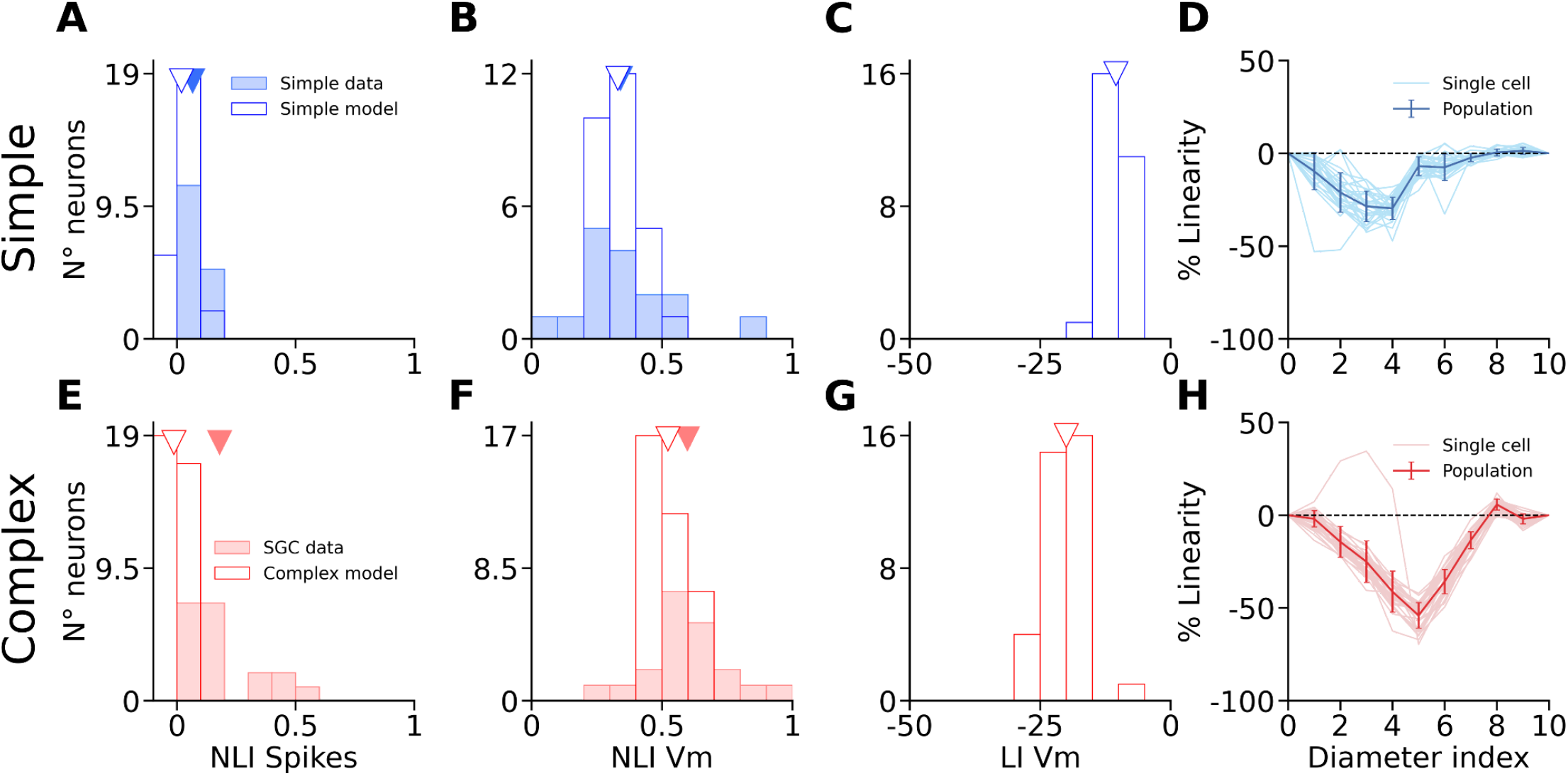
Distributions of non-linearity metrics in the model. **A**. Distribution of NLIs computed from F1 spikes for both data simple cells (filled blue bars) and model simple cells (hollow blue bars). Triangles mark the average for each population of the corresponding color. **B**. Same as **A** with NLIs computed from VmF1. **C**. Same, with LI computed from VmF1. **D**. % linearity for VmF1 in individual simple cells (pale blue) and population average across all model simple cells (blue) for each disk-ring pair. **E**. Distribution of NLIs computed from the mean firing rates for both data SGC (filled red bars) and model complex cells (hollow red bars). Triangles mark the average for each population of the corresponding color. **F**. Same as **E** with NLIs computed from VmDC. **G**. Same, with LI computed from VmDC. **H**. % linearity for VmDC in individual complex cells (pale red) and population average across all model complex cells (red) for each disk-ring pair.

We also calculated the Linearity Index (LI) of the membrane potential and found that the mean value for complex cells (LI VmDC = -20.0 ± 3.86) was lower than that of simple cells (LI VmF1 = -10.4 ± 2.34) (Fig. 8CG), with the difference being statistically significant between the two groups (Simple cells median: -10.7, Complex cells median: -20.1, p < 0.0001, U=978, Mann-Whitney), in line with cats (Cat simple cells median: -22.6, cat SGC median: -52.4, p = 0.0002, U=279, Mann-Whitney). Furthermore, the relative profiles of LIs for each diameter value is very similar to that of the data. In both data (Figure 6A) and model (Figure 8DH), simple cells exhibit a peak LI for smaller diameter values and with smaller magnitude than complex cells. Overall, we however observe lower magnitudes of LIs in the model than in cat data. This is due to the aforementioned weak suppression in the Vm of model neurons in response to large disks. As the denominator in the LI equation represents the response to the largest disk, the stronger suppression in cat cells increases the absolute magnitude of LIs in the cat relative to model neurons.

We have shown that our model reproduces the linear vs. non-linear spatial integration properties of simple vs. SGC cells. Interestingly the IGC neurons have similar basic connectivity principles as SGC cells: they send long-range lateral connectivity and receive weak direct thalamic input unlike the granular layers, yet the experiments revealed that the spatial integration properties of IGC cells are significantly more linear than SGC cells. To understand how such different spatial integration properties emerge, despite similar connectivity properties of SGC vs IGC, we have systematically investigated parameters of the layer 2/3 connectivity which is responsible for the non-linearity of spatial integration in this model layer. We found that parameters influencing the balance of long-range excitatory vs. inhibitory interactions can explain the shift in spatial integration linearity. Specifically, varying the spatial range or the iso-orientation bias of the horizontal connections targeting excitatory neurons, while keeping the parameters of those targeting inhibitory neurons fixed (Fig. 9A), has a strong impact on the spatial integration properties of the model layer 2/3 neurons.

**Fig. 9:**
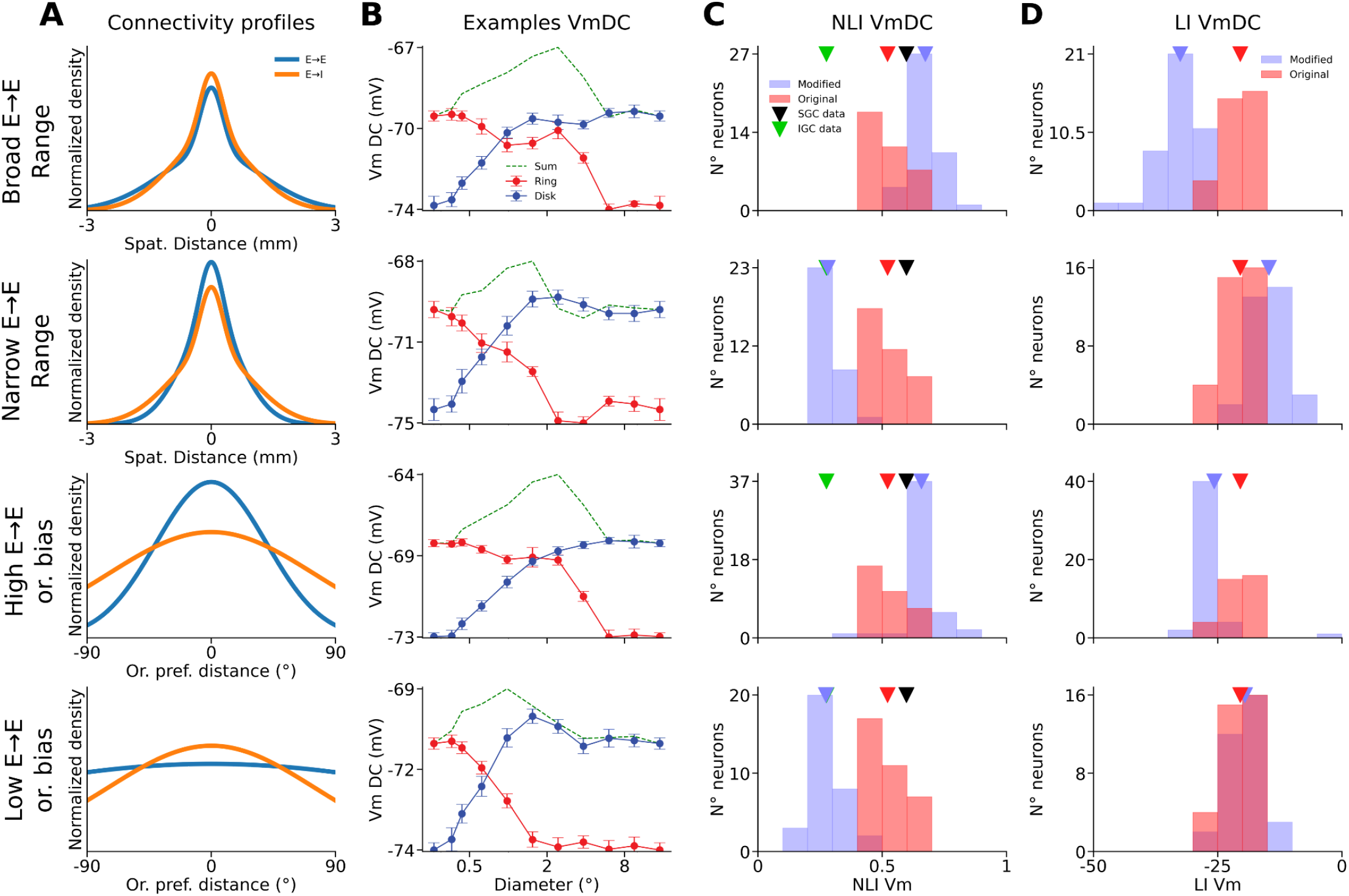
Impact of model parameters on complex cells linearity. **A**. Representations of the four modified configurations of the model (see Methods) based on the probability distribution of the recurrent connectivity depending on spatial distance between pairs of neurons in layer 2/3 (top two rows), or based on the probability distribution of the long-range component of the recurrent connectivity depending on orientation preference difference between pairs of neurons in layer 2/3 (bottom two rows). First row: Excitatory-to-excitatory connections (blue) are spatially broader than excitatory-to-inhibitory connections (orange). Second row: Excitatory-to-excitatory connections are spatially narrower than excitatory-to-inhibitory connections. Third row: Excitatory-to-excitatory connections are more tuned to orientation than excitatory-to-inhibitory connections. Fourth row: Excitatory-to-excitatory connections are less tuned to orientation than excitatory-to-inhibitory connections. **B**. VmDC of a model layer 2/3 complex cell in response to disks (blue) and rings (red) for each model configuration. In all cases the baseline has been subtracted. The green dashed lines represent the sum of the response to both stimuli in respect to baseline. **C**. Distribution of NLIs computed from VmDC of layer 2/3 complex cells for both the original version of the model (red bars) and each new configuration (blue bars). Triangles mark the averages for the original model (red), for each new configuration (blue), for data SGC (black) and for data IGC (grey). **D**. Same as **B**, but for LI instead of NLI.

Increasing the spatial range or the orientation bias of the long-range connections targeting excitatory neurons increases Vm depolarization in layer 2/3 complex cells for a wide range of inner diameters of ring stimuli, as can be observed in the example cells in Fig. 9B (first and third rows). In these cells, both disks and rings with diameters and inner diameters of 2.4° elicit a very strong depolarization relatively to the peak depolarization observed, making them behave even more non-linearly than in the example shown from the original model (Fig. 7F). The NLI of VmDC of the cells is therefore shifted towards higher values (broader spatial range: 0.67 ± 0.06, higher iso-orientation bias: 0.67 ± 0.05) (Fig. 9C, first and third rows). Inversely, decreasing either the spatial range or the orientation bias of the long-range connections targeting excitatory neurons induces a much steeper drop of the depolarization elicited by ring stimuli as their inner diameters increase (Fig. 9B, second and fourth rows). Therefore, there is never a pair of disks and rings stimuli of similar outer and inner diameters respectively that both elicit a significant depolarization, resulting in a decrease of the NLI of the Vm of layer 2/3 complex cells, to a level similar to that of cat IGCs (narrower spatial range: 0.28 ± 0.05, lower iso-orientation bias: 0.27 ± 0.07) (Fig. 9C, second and fourth rows). As expected, parameter changes had the inverse effect on LIs (broader spatial range: -32.6 ± 3.91, narrower spatial range: -14.7 ± 3.06, higher iso-orientation bias: -25.7 ± 6.24, lower iso-orientation bias: -19.5 ± 3.23) (Fig. 9D).

## Discussion

Our results reveal an important difference in the spatial integration properties between complex cells in supra and infragranular layers of the primary visual cortex of the cat. Using complementary pairs of circular (“disks”) and annuli (“rings”) patches of optimal drifting gratings, we showed that: (i) Rings with inner diameter equal or greater than the diameter of disk eliciting maximal response, still triggered a strong depolarizing input (VmDC) and strong postsynaptic activity as estimated through VmSD in supragranular complex cells but almost none in infragranular complex cells or in simple cells (see Fig 4). (ii) The ratio of the maximum predicted divided by the maximum obtained response, i.e. the non-linearity index (NLI), was greater for SGC than for IGC cells. (iii) The ratio of the sum of each disk-ring separate responses divided by the response to them presented together, i.e., the linearity index (LI), showed that SGC neurons perform highly sublinear summation of synaptic input while IGC cells are close to linear. Simple cells showed a small degree of sublinearity. (iv) Using a recurrent spiking model of cat V1 we have demonstrated that the differences between supra and infragranular layers can be explained by different patterns of long range horizontal connectivity onto excitatory and inhibitory cells of the local networks.

### Suppression

In almost all cells, regardless of location, we observed an increase in response amplitude to a maximum followed by some degree of suppression. Response suppression has been extensively studied using extracellular recordings (Hubel and Wiesel, 1962; Bolz and Gilbert, 1986; DeAngelis et al., 1992, 1994; Gilbert et al., 1996; Sengpiel et al., 1997; Walker et al., 2000) and less so intracellularly (Ozeki et al., 2009). It is important to emphasize that we only use one stimulus orientation and therefore the phenomenon we report here is similar to end stopping, or what was originally called hypercomplex cells by Hubel and Wiesel (1962). The first consideration is whether this form of suppression is caused by an increase in inhibition. When activated alone, the input from the region that causes suppression not only depolarized SGC cells but also increased the amount of synaptic input as measured by the standard deviation of the Vm (the VmSD). This is contrary to the presence of inhibition which by definition, when elicited close to its reversal potential, strongly reduces the input resistance, effectively “shunting” integration of other synaptic input. In other words, if the depolarization was due to our inability to detect inhibition because of the low resting Vm, its presence would have been revealed by a strong reduction in VmSD, but the VmSD decreased only in parallel with the VmDC. Thus, our results do not support a strong engagement of inhibition. A similar conclusion was reached with iontophoresis experiments using extracellular recordings in cat V1 (Ozeki et al., 2004). Also in agreement is the intracellular study of Ozeki et al. (2009). They showed that during iso-orientation surround suppression both excitatory and inhibitory conductances estimated from the Vm are reduced and this effect is intracortical, i.e., not due to reduction in LGN input. This result concurs with the length tuning results of Anderson et al. (2001). The study of Ozeki et al. (2009) provided strong evidence that during visual stimulation, cat visual cortex operates as an inhibition-stabilized network (ISN), a type of paradoxical network effect (as first shown by Tsodyks et al., 1997), where inhibitory neurons response overshoots that of excitatory neurons in response to a strong external excitatory input. This transient increase in inhibition leads to a decrease of the recurrent input from the local excitatory cells, resulting in a withdrawal of both excitation and inhibition within the local network. Using a computational model, Ozeki et al. (2009), showed that an ISN provides the best explanation for the network behavior underlying response suppression by the iso-orientation surround. The ISN regime is compatible with the behaviour of SGC cells revealed in our study, predicting both the presence of net excitatory input from the surround, as revealed by our ring experiments, and the net inhibitory impact on Vm and spiking response of neurons when the matching pair of disk and ring stimuli is presented simultaneously. However, our findings open a new question of how can the V1 circuitry support both the linear and non-linear integration regimes and why do they systematically vary across cortical layers, which we address in the multi-layer model presented in this study.

### Residuals

Our measurements of synaptic input triggered from the area of visual space beyond the diameter of maximal response show that SGC cells have a strong depolarizing input but IGC cells do not. This difference between SGC and IGC cells in spatial integration revealed by the residuals, was only at the level of synaptic responses and did not translate into their spike output. The spike output residual was not statistically different between complex cells at different depths. It is possible that the much smaller number of spikes collected during intracellular recordings, due to their shorter recording time compared to extracellular recordings, was not sufficient to capture the effect. The presence of positive residual values in the spike output of some SGC cells (as shown in Fig. 4I) may be suggestive of such an undersampling problem. However, it is also possible that, indeed, the phenomenology described here occurs only at subthreshold Vms. This begs the question of its functional significance. We propose that the high synaptic residual values observed in SGC but not IGC cells are critical in second order interactions between visual input. Indeed, when the ring is presented together with the disk it causes response suppression (see Fig. 4A), but when presented alone it causes excitation without any evidence of strong inhibition (see Fig. 4L). This shows that the same region of visual space generates two types of responses depending on whether the region occupied by the disk is stimulated or not. It is suggestive of complex interactions at the circuit level that lead to response normalization (Heeger, 1992; Carandini et al., 1997; Ozeki et al., 2004, 2009; Cardin et al., 2008; Carandini and Heeger, 2012). Such interaction is highly dependent on the relative properties of the two regions of space, such as contrast, orientation and so on and likely to play an important role in the integration of visual input of complex, or naturalistic, visual stimuli (Fregnac et al., 1996; Bringuier et al., 1999b; Ozeki et al., 2004; Angelucci and Shushruth, 2013; Gerard-Mercier et al., 2016). Experiments in course in our lab suggest that to be the case.

### Non-linearity index

The non-linearity index (NLI) is a comparison of the maximum algebraic sum with the maximum obtained response. It compares the maximum predicted response of a cell performing linear summation, with the maximum response it actually generated. The NLI then can be understood as a simple and global measure of non-linearity. Very importantly, the NLI does not take into consideration spatial boundaries of center or surround and is calculated for the entirety of the visual area that triggers any synaptic response. The NLI is internally consistent as the prediction of maximum linear response for each cell is based on the synaptic responses of that cell, rather than an external model. Our results showed that using this metric distinguishes SGC and IGC cells. It also shows that ALL our cells including simple cells perform sublinear summations.

### Linear summation

Integration of inputs within the RF of simple and complex cells has been studied with intracellular recordings. Ferster (1994) have shown that simple cells sum synaptic inputs linearly *within* their RFs by demonstrating that the response to drifting gratings can be accurately predicted by applying the responses to stationary stimuli to a purely linear model (Jagadeesh et al., 1993). In contrast, the presentation of brief bars within the RF of simple cells revealed a strong supralinearity when the two bars were presented within 10 ms of each other (Cardin et al., 2010). Intracellular recordings from complex cells demonstrated strong sublinearities of summation of synaptic potentials from oriented bars presented within the RF (Lampl et al., 2004). Here we measured linearity by comparing pairs of stimuli that, when presented together, encompass the entirety of the region triggering synaptic responses in each neuron. We observed that SGC cells are on average highly sublinear but that the sublinearity is stronger for middle values of diameter (see Fig. 6A), i.e., when the disk and ring responses become comparable in size. Thus, integration properties will vary in consonance with varying stimulus contrast distributions, such as during natural vision.

There are many sources of non-linearities in neurons (Koch, 1998). These include intrinsic currents (Llinas, 1988), shunting inhibition (Fatt and Katz, 1953; Eccles, 1964), short term synaptic plasticity (Zucker and Regehr, 2002), network effects and many others. For example, sublinearity of synaptic input summation may result from physical proximity of excitatory inputs in the dendrite. Physical proximity leads to shunting of their postsynaptic conductances with a corresponding decrease in the generation of postsynaptic voltage and thus, sublinear summation (Bernander et al., 1991). In contrast, similar inputs in different dendrites will add up linearly when their electrotonically propagated potentials reach the soma (Koch, 1998). This mechanism is relevant because the SGC pyramidal dendritic arbors are much smaller than IGC pyramidal cells and may therefore lead to a pronounced shunting. Exploring these possibilities would require detailed multicompartmental models.

### Circuits

In the neocortex, the laminar separation of neurons implies different positions in local and long circuits. With respect to our work, supragranular pyramidal cells in sensory cortices receive a major driving input from layer 4 and send output to other cortical columns in the form of horizontal unmyelinated axons (as well as myelinated axons to homologous contralateral cortex). Infragranular layer cells represent the output of the neocortex to subcortical structures. With the exception of the upper tier L6 simple cells, which are functionally driven by LGN input and in return project mainly to LGN, the rest of infragranular cells are functionally complex cells and receive their input from their neighbors, from supragranular cells and from other cortical areas. Thus, SGC and IGC cells represent two very different circuit nodes, SGC sending output to cortex for further processing of image attributes, the other sending output to brainstem for systems related to action, such as cortico-pontocerebellar fibers involved in the coordination of visually guided movement as well as coordination of eye movements (Baker et al., 1976), or fibers to superior colliculus (Rosenquist and Palmer, 1971, 1971), which is the origin of the tectospinal tract for orienting movements of head and eyes towards moving stimuli. We posit that nonlinear summation is part of the computations required for processing of visual image, while linear summation is a rapid form of faithfully relaying visual input for action.

### Horizontal connections

Cortical columns are interconnected by a vast system of horizontal axons from pyramidal cells in supragranular layers that terminate in small clusters of synaptic buttons roughly the size of a column (Ts’o et al., 1986; Gilbert and Wiesel, 1989; Hata et al., 1991; Bosking et al., 1997; Angelucci and Bullier, 2003; Martin et al., 2014, 2017; Chavane et al., 2022). These buttons terminate in both excitatory and inhibitory neurons in the local circuit within the column (Binzegger et al., 2004). The synaptic strength of the terminals is determined by the number of buttons per cell, the probability of release, vesicle size, electrotonic position in the dendritic tree of the postsynaptic cell and perhaps, more importantly, the degree of synchronization of the input. Despite a vast knowledge of horizontal connection terminals in supragranular layers 2-3, almost none of the elements discussed above is known. Our model suggests a plausible and powerful mechanism based on horizontal connectivity to explain the difference between SGC and IGC cells

### Computational model

We present a computational model of layers 4 and 2/3 of V1 deeply anchored in architecture of known V1 circuitry and function (Antolík et al., 2024), which reproduces the differences in spatial integration between simple cells and SGC cells. In both model simple and complex cells, firing rates exhibit linear spatial input integration, whereas VmDC of complex cells display significant sublinearity. We found that increasing the spatial range of long-range connections targeting excitatory cells or increasing the iso-orientation bias of those connections increased the sublinearity of layer 2/3 cells. Conversely, decreasing the values of those parameters led to a decrease of sublinearity to a level similar to observed experimentally in IGC.

Therefore, our modeling results suggest that linearity of spatial integration of primary visual cortical complex cells may depend on the spatial and functional properties of long-range excitatory horizontal connections. For a given orientation column, long-range excitatory connections drive both direct excitatory and indirect inhibitory (via local interneurons) inputs. When horizontal connections targeting excitatory cells reach laterally further or exhibit higher iso-orientation bias relative to the long-range excitatory connections targeting inhibitory cells, the balance of Excitation/Inhibition from long-range inputs shifts towards net excitation (Fig. 10A). This leads to strong Vm depolarization originating from the area beyond the maximum response when this area is stimulated in isolation. However, in response to a large disk, the presence of a strong feedforward drive in addition to the horizontal input makes their membrane response saturate due to the higher gain of inhibitory compared to excitatory cells (Fig. 10C), in line with an inhibitory stabilized regime (Tsodyks et al., 1997; Ozeki et al., 2009). Inversely, when long-range excitatory-to-excitatory connections have a narrower spatial range or are less tuned than long-range excitatory-to-inhibitory connections, contribution of the stimulation of the surround shifts towards being balanced or net inhibitory (Fig. 10B). This leads to a rapid decline in the Vm response of the cells as the inner diameter of the ring stimuli increases and stimulus withdraws from visual space mediated by thalamo-cortical connections to the given neuron. Our data did not show an overt role for inhibition triggered from the rings that causes suppression, favoring the balanced influence of surround input in IGCs. In order to test this hypothesis, experiments need to couple the visual stimuli with current injection in order to estimate excitatory and inhibitory conductances in supra- vs. infra-granular layers. Furthermore, experiments must distinguish between excitatory and inhibitory cells in order to test the connectivity hypothesis proposed by our model. Note that our simulations predict that even small magnitude of such functional differences between excitatory-to-excitatory and excitatory-to-inhibitory long-range connections can have major consequences for spatial integration of sensory inputs in the cortex and may explain the difference in spatial integration properties between IGC and SGC cell populations discovered by our data.

**Fig. 10:**
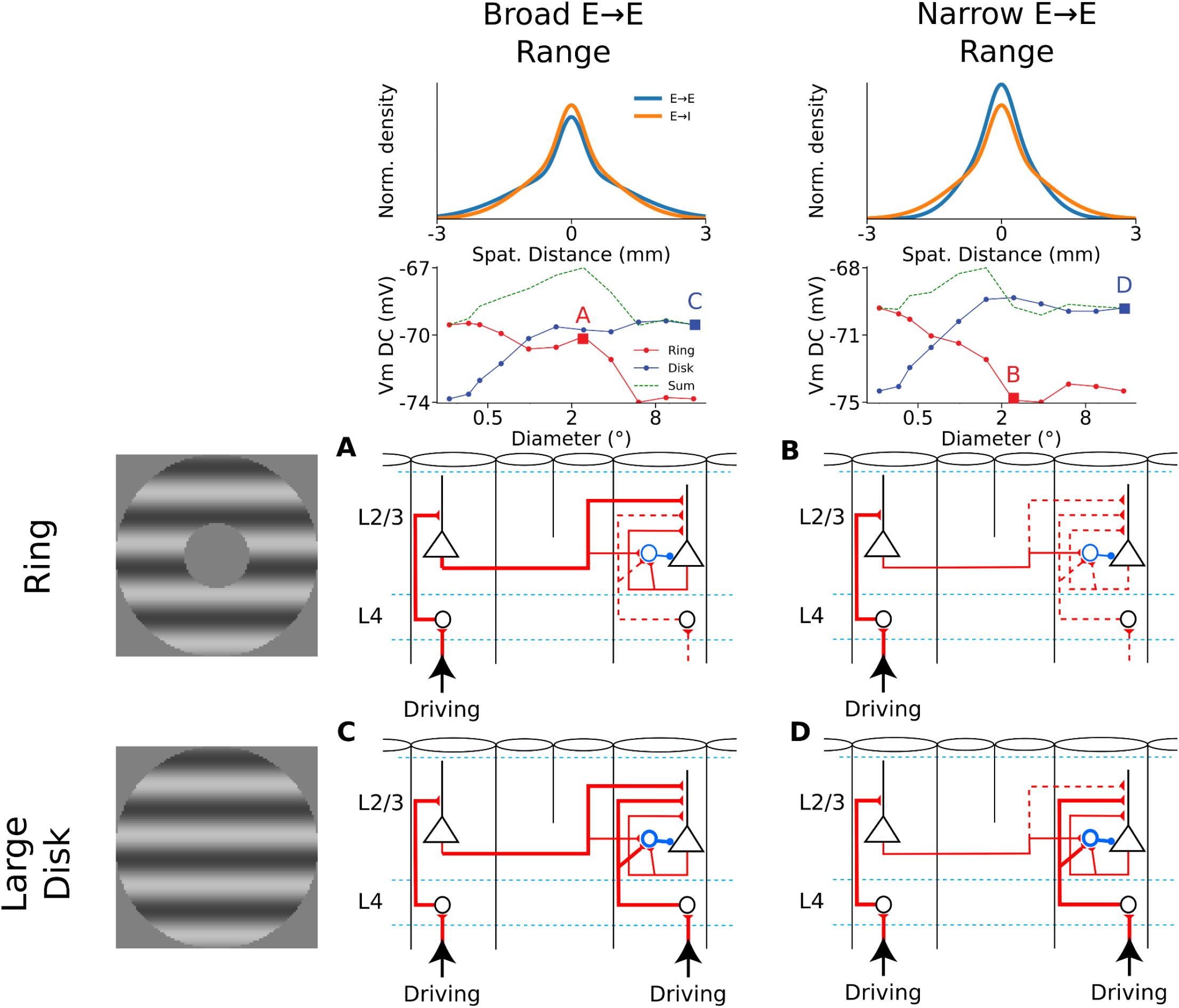
Impact of model parameters on complex cells linearity. **A**. Mechanistic explanation of the response of the model configuration with spatially broad excitatory-to-excitatory connections to a ring with a hole covering the center of the stimulus. **B**. Same as **A** for the model configuration with spatially narrow excitatory-to-excitatory connections. **C**. Same as **A** but in response to a large disk. **D**. Same as **B** but in response to a large disk.

We chose to study the mechanisms that control the non-linearity in the model SGC cells, instead of explicitly modelling the infra-granular layers, due to the lack of accurate quantitative analysis of the physiology and circuit properties of the deep layers. The gross intra-V1 anatomical features of the supra- and infra-granular layers are very similar, both receiving primarily inputs from other V1 layers, and both sending long-range V1 connections. However, at finer detail and unlike in supra-granular layers, the anatomical and functional biases of the infra-granular connectivity is very poorly characterised, leaving us with little to differentiate a hypothetical explicit model of infra-granular layers. Yet, as our results show, it is exactly such properties that have a major impact on spatial integration. Another limitation of our model is the absence of feedback from higher visual areas in the model, which have been shown to participate in size tuning of V1 cells (Nurminen et al., 2018). The lack of inputs from higher visual areas may thus be the cause of the weaker surround suppression of Vm in model cells relative to cat data, which in turn leads to lower LI in the model in comparison to the experimental data. Finally, each layer of our model is composed by only one population of inhibitory cells and thus cannot account for potential differential roles played in spatial input integration by parvalbumin or somatostatin expressing cells (Adesnik et al., 2012) in the phenomena presented here.

We are not aware of any other V1 modeling studies which investigated the response of a large-scale spiking neural network to ring stimuli. The model presented here is thereby the first model instance to reproduce accurately the level of non-linearity in spatial integration of both granular and supragranular layers which was brought to light through this experimental protocol. However, models comparable to ours have been used to study spatial integration in V1 with disk stimuli. Somers et al. have shown in a spiking neural network with functionally biased connectivity how realistic contrast and orientation dependent surround suppression can be obtained via having higher input gain in inhibitory neurons than in excitatory neurons (Somers, 1998). The modeling series from Wielaard et al. also reproduces size tuning in a single instance of V1 model which also exhibits orientation tuning, spatial and temporal frequencies tuning as well as the emergence of a bimodal distribution of simple and complex cells (Wielaard and Sajda, 2006a, 2006b).

## Additional information

### Author Contributions

D. C. and L. P. conceived the experimental protocol and recorded the experimental data. D. C., R. C. and J. A. designed and performed the data analysis. R. C. designed the computational model and ran the computational simulations under the supervision of J. A.. D. C. and R. C. created the figures. The four authors jointly wrote and reviewed the manuscript.

### Data availability

All Intracellular recordings are stored in Neuralynx format or Igor Pro 9.0 and are available upon request. Routines for data analysis of experimental data were written in Igor Pro and are also available upon request.

The V1 model, together with the simulation experiments and analyses, was implemented through the Mozaik framework (Antolík and Davison, 2013), freely accessible at https://github.com/CSNG-MFF/mozaik (version 0.5). To run the model one first needs to install Mozaik and its dependencies as indicated in the Mozaik Github repository, and then to download the implementation of model at https://github.com/CSNG-MFF/mozaik-models/tree/main/Cagnol2025 (version 1.1) and to follow the instructions to run the experiments presented in this study. We used NEST simulator (version 3.4) as the underlying simulation engine (Sinha et al., 2023).

